# Rapid endothelial infection, endothelialitis and vascular damage characterise SARS-CoV-2 infection in a human lung-on-chip model

**DOI:** 10.1101/2020.08.10.243220

**Authors:** Vivek V Thacker, Kunal Sharma, Neeraj Dhar, Gian-Filippo Mancini, Jessica Sordet-Dessimoz, John D McKinney

## Abstract

Severe cases of COVID-19 present with hypercoagulopathies and systemic endothelialitis of the lung microvasculature. The dynamics of vascular damage, and whether it is a direct consequence of endothelial infection or an indirect consequence of immune cell mediated cytokine storms is unknown. This is in part because *in vitro* models are typically epithelial cell monocultures or fail to recapitulate vascular physiology. We use a vascularised lung-on-chip model where, consistent with monoculture reports, low numbers of SARS-CoV-2 virions are released apically from alveolar epithelial cells. However, rapid infection of the underlying endothelial layer leads to the generation of clusters of endothelial cells with low or no CD31 expression, a progressive loss of endothelial barrier integrity, and a pro-coagulatory microenvironment. These morphological changes do not occur if these cells are exposed to the virus apically. Viral RNA persists in individual cells, which generates a response that is skewed towards NF-KB mediated inflammation, is typified by IL-6 secretion even in the absence of immune cells, and is transient in epithelial cells but persistent in endothelial cells. Perfusion with Tocilizumab, an inhibitor of trans IL-6 signalling slows the loss of barrier integrity but does not prevent the formation of endothelial cell clusters with reduced CD31 expression. SARS-CoV-2 mediated endothelial cell damage occurs despite a lack of rapid viral replication, in a cell-type specific manner and independently of immune-cell mediated cytokine storms, whose effect would only exacerbate the damage.

## Introduction

Organ-on-chip technologies recreate key aspects of human physiology in a bottom-up and modular manner^1^. In the context of infectious diseases, this allows for studies of cell dynamics^2,3^, infection tropism^4^, and the role of physiological factors in disease pathogenesis in more native settings^5^. This is particularly relevant for the study of respiratory infectious diseases^6^, where the vast surface area of the alveoli poses a challenge to direct experimental observation.

COVID-19, caused by the novel *betacoronavirus* SARS-CoV-2, first manifests as an infection of the upper airways. Severe cases are marked by progression into the lower airways and alveoli. Here, it manifests as an atypical form of acute respiratory distress syndrome (ARDS) characterized by good lung compliance measurements^7,8^, and elevated levels of coagulation markers such as D-dimers^9^, and pro-inflammatory markers in the blood^10^. Autopsy reports show numerous microvascular thrombi in the lungs of deceased patients together with evidence of the intracellular presence of the virus in vascular cells^11,12^. These reports suggest that infection of and alterations to lung microvasculature plays a key role in COVID-19 pathogenesis^13,14^, which is further reinforced by the early observation of endothelial cell damage, platelet aggregation and the formation of microthrombi in an ante-mortem study^15^. Yet most *in vitro* studies have focused on monocultures of upper airway respiratory cells. In studies with alveolar epithelial cells, SARS-CoV-2 has been shown to replicate poorly both in the A549 lung adenocarcinoma cell line^16^ and in primary alveolar epithelial cells *ex vivo*^17^ and has been reported to be unable to infect primary lung microvascular endothelial cells^18^, which are at odds with the reported medical literature. There is therefore an urgent need for a better understanding of the pathogenesis of SARS-CoV-2 in alveolar epithelial cells, especially in a more realistic model of the alveolar space that is vascularized. The lung-on-chip model is well-suited to this purpose because it includes a vascular compartment maintained under flow, and infection can occur at the air-liquid interface^19^, two key physiological features that are lacking in organoid models^20–22^. We therefore establish a human lung-on-chip model for SARS-CoV-2 infections, and probe the viral growth kinetics, cellular localization and responses to a low dose infection using qRT-PCR, ELISA, RNAScope, immunofluorescence and confocal imaging (Fig. S1A).

## Results

### SARS-CoV-2 infection of a human lung-on-chip alters expression of viral entry factors

A human lung-on-chip (LoC) model for SARS-CoV-2 pathogenesis (Fig. 1A) mimics the alveolar space using primary human alveolar epithelial cells (‘epithelial’) and lung microvascular endothelial cells (‘endothelial’), which form confluent monolayers on the apical and vascular sides of the porous membrane in the chip (Fig. 1B, C). The modular nature of the technology allowed us to recreate otherwise identical LoCs either without (‘w/o’) (Fig. 1B) or with the addition of CD14+ macrophages (representative image in Fig. S1B) to the epithelial layer on the apical side of the chip. The latter configuration was used for a subset of experiments as indicated throughout the text. We first characterized the epithelial and endothelial cells used, both in monoculture and on-chip at the air-liquid interface to verify that the cells mimicked human alveolar physiology. In epithelial cells, *ACE2* expression was low both in monoculture (Fig S2A) and on-chip (Fig. 1E) consistent with transcriptomic^18,23^ and proteomic analyses^24^ of human tissue. Expression on-chip was five-fold lower than in monoculture (Fig. S2B). Low *ACE2* expression was also observed from RNAscope assays on uninfected LoCs (Fig. 1D). Although *ACE2* expression was particularly low in endothelial cells in monoculture (Fig. S2A, expression increased 10-fold on-chip (Fig. S2B) so that levels in epithelial and endothelial cells were comparable (Fig. 1E). This upregulation in *ACE2* expression could possibly be due to the effects of shear stress and flow in the vascular channel^25^. Neuropilin-1, an integrin-binding protein^26^ has recently been reported to be an alternative receptor for SARS-CoV-2 entry^27,28^ that is highly expressed in pulmonary microvascular cells. Consistent with these reports, *NRP1* expression was between one and four orders of magnitude higher than *ACE2* expression in both cell types in monoculture (Fig. S2A) and on-chip (Fig. 1E). On-chip, *NRP1* expression was 6-fold higher in endothelial cells compared to epithelial cells (Fig S2B). Lastly, mature alveolar epithelial cells retained differentiation in type I and type II cells, which we verified by immunostaining for type II (pro-SPC) and type I (Podoplanin) markers in monocultures (Fig. S3A-C) as well as through qRT-PCR measurements for type II (*ABCA3*, *SFTPC*) and type I (*AQP5*, *PDPN*, *CAV1*) markers in cells from the epithelial layer of uninfected LoCs (Fig. S3D). Endothelial cells showed characteristic expression of Platelet Endothelial Cell Adhesion Molecule (PECAM-1 or CD31) at cell junctions (Fig. 1C, additional images in Fig. 4C, D).

**Fig. 1.**
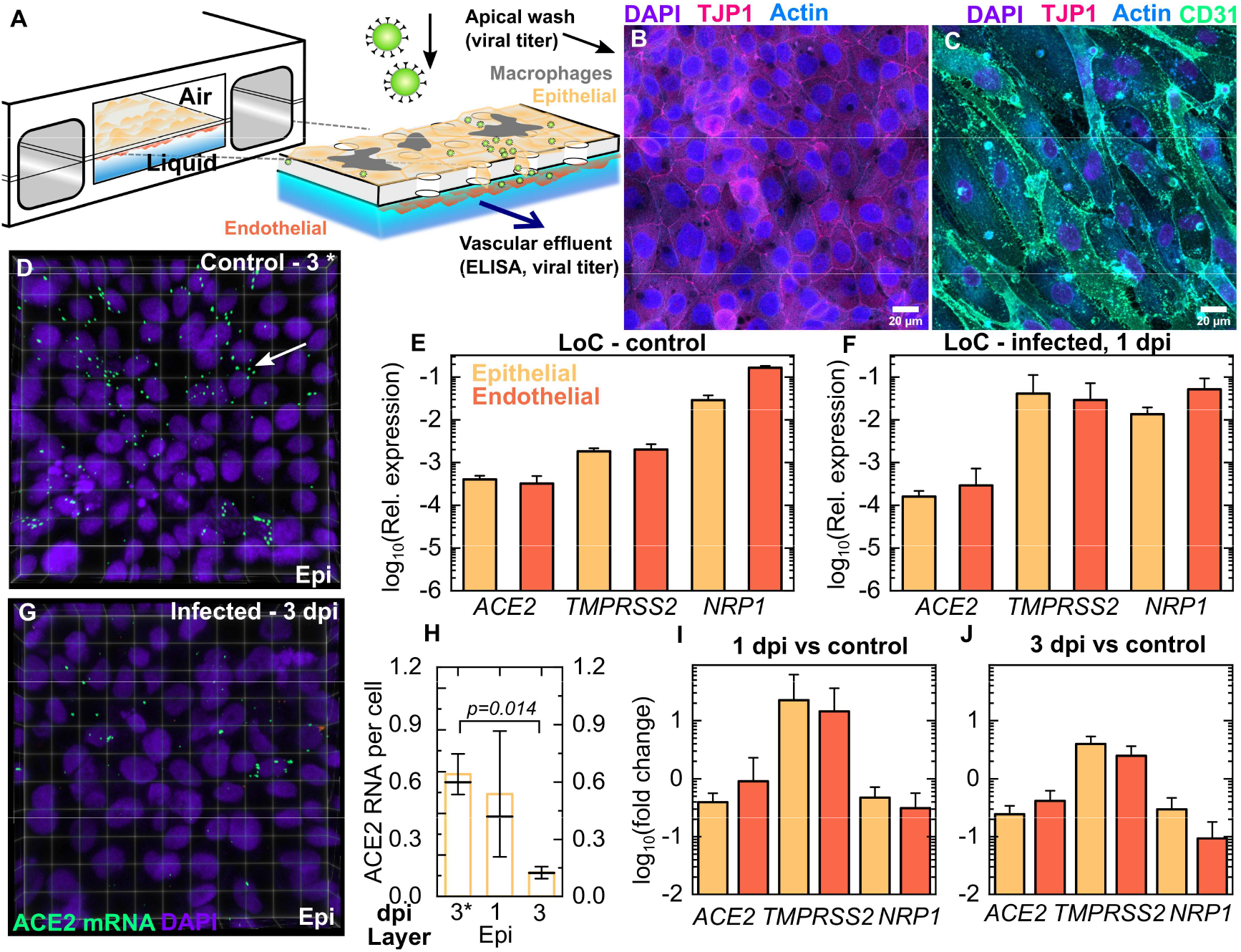
SARS-CoV-2 infection in a human lung-on-chip model. (**A**) Schematic of the LoC model for SARS-CoV-2. Maximum intensity projections show that confluent layers of epithelial and endothelial cells with strong expression of tight junction markers populate the top (**B**) and bottom faces (**C**) of the porous membrane. Actin, CD31, TJP-1 and nuclear labelling are shown in azure, spring green, bright pink, and electric indigo LUTs respectively. qRT-PCR characterization of the epithelial and endothelial layers on-chip for SARS-CoV-2 entry markers (*ACE2, TMPRSS2, NRP1*) for cells extracted from the apical and vascular channels from n=3 uninfected LoCs (**E**) and n=4 infected LoCs at 1 day-post infection (**F**). In both cases expression is normalized to *GAPDH* levels. *ACE2* levels were also characterized using RNAscope, a representative 3D view of a 232 × 232 μm^2^ field of view of the epithelial face of an uninfected control at 3 days at air-liquid interface (3C) (**D**) and infected LoC at 3 dpi (**G**) are shown. The nuclear labelling is indicated in electric indigo and *ACE2* mRNA is labelled in spring green LUT. (**H**) *ACE2* expression decreases between 1and 3 dpi in the LoC model, quantified on a per-cell basis for 4-5 fields of view across one control and one infected chip at each timepoint. (**I, J**) A plot of the fold change in expression of host proteins required for viral entry at 1 day-post infection (n=4 LoCs) and 3 days-post infection (n=2 LoCs) vs. uninfected controls (n=3 LoCs). The bars represent the mean value, the solid line represents the median value, and the error bars represent the standard deviation. P-values are calculated using a Kruskal-Wallis one-way ANOVA test.

A low dose infection (200-300 plaque forming units (PFUs), multiplicity of infection (MOI)=0.003) of the apical surface of the LoC resulted in decreases in *ACE2* expression in epithelial and *NRP1* expression in both epithelial and endothelial cells at 1 dpi (Fig. 1F, I). Expression of both cell surface receptors remained downregulated at 3 dpi (Fig. 1J). These results are consistent with observations using RNAscope assays, where *ACE2* levels in the epithelial layer declined over subsequent days (Fig. 1D, G, H, p=0.014). On the other hand, infection resulted in increased expression of the TMPRSS2 protease^29^ required to cleave the spike protein for viral entry at 1 dpi in both cell types relative to uninfected controls (Fig. 1 F, I), with variable expression levels across chips. This trend was also maintained at 3 dpi (Fig. 1J). Overall, these results show that both ACE2 and NRP1 likely play a role in pathogenesis in the alveolar space, aided by increased TMPRSS2 levels. In comparison, infection did not alter levels of type II or type I alveolar epithelial cell markers significantly (Fig. S3E, F).

### Infection of the alveolar space is characterized by a lack of productive infection, slow intracellular replication, and transmission to the endothelial layer

We first characterized the progression of infection by measuring the release of viral genomes via qRT-PCR and the number of infectious virions released via measurement of plaque forming units (PFUs). Infected LoCs were monitored daily for the release of infected viral progeny (1) *apically* – on the epithelial layer and (2) in the cell culture media flowed through the vascular channel (‘apical wash’ and ‘vascular effluent’ in Fig. 1A). A low number of viral genomes were released apically from the epithelial layer, and the number of genomes detected decreased over 1-3 dpi (Fig. 2A). Genome copy numbers were 100-fold lower than the starting inoculum (between 200-300 PFU). Low numbers of infectious virions were also detected via measurements of PFU from eight LoCs w/o macrophages at 1 dpi and two LoCs w/o macrophages at 2 dpi (Fig. 2B). These results showed that, on the whole, productive infection of the epithelial cell layer did not occur on-chip, although instances of small numbers of epithelial cells with high levels of spike protein (Fig. S5A) or viral genomic RNA (Fig. S9) were detected via microscopy and likely contribute to the low levels of infectious virions released. No viral genomes were detected in the vascular effluent (Fig. 2B), and the lack of infectious particles in the effluent was confirmed for two LoCs with and w/o macrophages by plaque forming unit assays (data not shown). This ruled out the possibility of direct release of virions from the epithelial cell layer into the vascular channel and suggested that either dissemination of virions to the endothelial cell layer did not occur, or that infection of the endothelial layer could occur but no viral genomes or infectious virions were released *apically* into the vascular channel by endothelial cells.

**Fig. 2.**
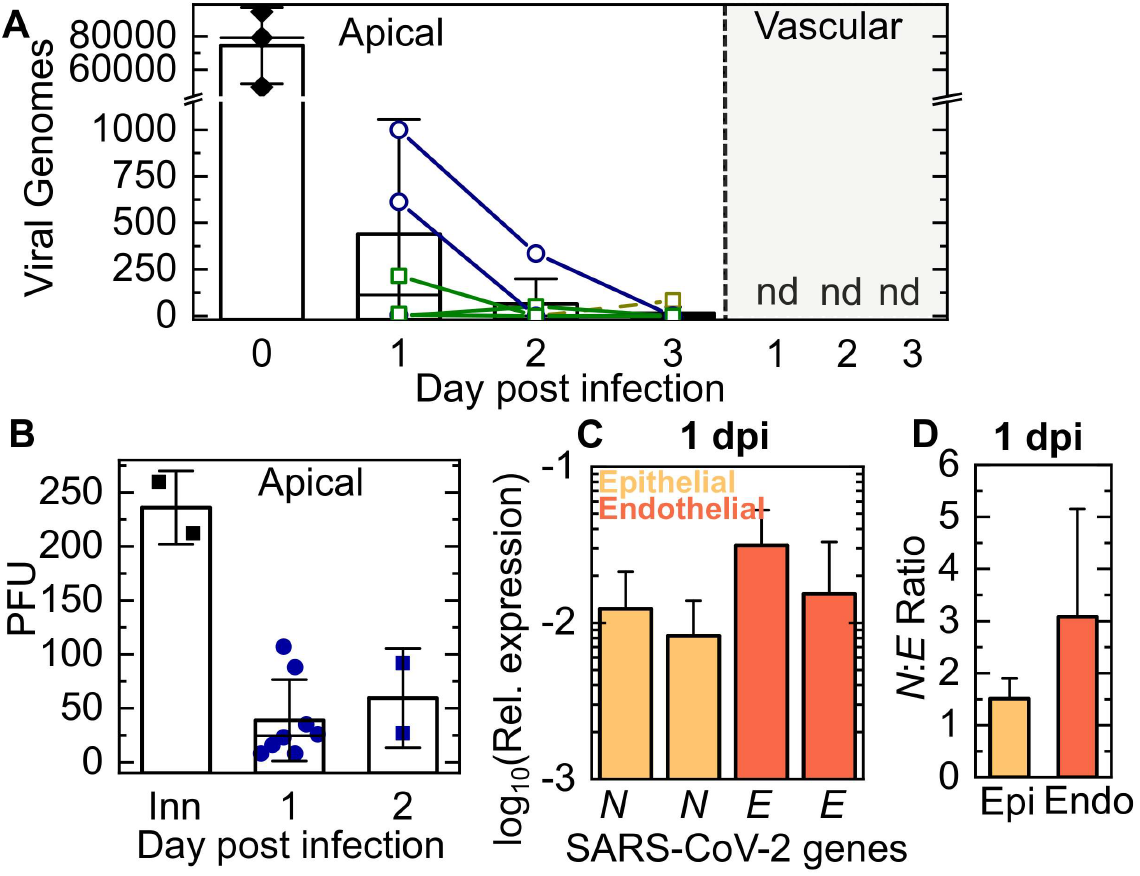
Viral replication but lack of release of infectious virions characterizes SARS-CoV-2 infection in the LoC model. The kinetics of release of SARS-CoV-2 progeny was assessed from samples obtained from a wash of the apical face of the chip (**A, B**), and the vascular effluent (**A**) collected at each day-post infection. (**A**) The number of viral genomes in each sample was quantified using a one-step qRT-PCR assay for the SARS-CoV-2 N gene. The starting inoculum introduced to the apical side (labelled ‘Inn’) corresponded to 200-300 PFU. Lines join datapoints from the same LoC, reconstituted with (green squares) and without (dark blue circles) macrophages. Viral genome numbers within the apical wash showed a declining trend over 3 days of infection. Viral genomes are not detected (‘nd’) in the vascular effluent for all the chips shown. (**B**) Measurements of plaque forming units (PFU) from the apical wash of LoCs reconstituted without macrophages (n=8 for 1 dpi, n=2 for 2 dpi) confirms the presence of infectious virions. (**C**) Quantification of the intracellular levels of transcripts for the SARS-CoV-2 *N* and SARS-CoV-2 *E* genes in the epithelial and endothelial layers of infected LoCs at 1 dpi (n=3) relative to expression of the eukaryotic housekeeping gene *GAPDH*. (**D**) SARS-CoV-2 *N*: SARS-CoV-2 *E* ratio in the epithelial and endothelial layers of the infected LoCs shown in (**C**). In all plots, the bars represent the mean value and the error bars represent the standard deviation, the solid black line in (**A**) represents the median.

To address this, we further quantified intracellular viral RNA loads from the total RNA extracted from cells of the apical and vascular channels of three infected LoCs w/o macrophages at 1 dpi. This revealed a significant number of viral transcripts in cells from both the epithelial and endothelial layers (Fig. 2C, Fig. S4C). Additional quantification of intracellular viral RNA from an infected LoC w/o macrophages using a SARS-CoV-2 specific one-step qRT-PCR kit revealed >10^4^ genomes in both epithelial and endothelial cells (Fig. S4A); genome copy numbers exceeded those for cellular housekeeping gene *RNAseP* (Fig. S4B). Similar numbers of intracellular viral genome copy numbers have been reported for infections of alveolar epithelial cells in monoculture^17^ and these numbers are modest in comparison to cells such as airway epithelial cells and Vero E6 cells where SARS-CoV-2 replicates productively ^30^. The difference in replication kinetics can also be appreciated from the fact that whereas the *N* gene is most abundant during productive infection in Vero E6 cells ^31^, we obtained a ratio of SARS-CoV-2 *N* to SARS-CoV-2 *E* <5 (Fig. 2D, Fig. S4D), consistent with the observation of low numbers of extracellular infection virions (Fig. 2B).

The detection of SARS-CoV-2 genomes in the endothelial layer within 1 dpi suggests rapid dissemination through *basolateral transfer* from the epithelial to the endothelial layer can occur. Although basolateral transmission has not been reported previously to be the significant mode of transmission for both SARS-CoV-2^32^ and SARS-CoV-1^33^ infections of monocultures of upper airway cells at the air-liquid interface, this could reflect aspects of infection unique to alveolar epithelial cells, or more generally, of infection of cells where productive infection is rare.

### SARS-CoV-2 infection of endothelial cells on-chip but not in monoculture alters cell morphologies

Recently published autopsy reports of COVID-19 patients revealed features of exudative alveolar damage^12^, and a rapid loss of alveolar epithelial cell barrier integrity could account for the basolateral transfer to endothelial cells. The LoC model is compatible with microscopy-based assays, which allowed us to assess changes in cellular physiology with high spatial resolution using confocal microscopy. Even at 3 dpi, the epithelial layer maintained a high degree of confluency (Fig, 3D, Fig. S6F), with only a few signs of cell death (Fig. 3D, white arrows). In stark contrast, staining for F-actin localisation revealed striking changes to the morphology of cells in the vascular channel. In comparison to the confluent layer of cells in an uninfected control LoC (3D view in Fig 3A, Fig. S6A) infected LoCs showed profound alterations in endothelial layer architecture (Fig 3B, C). At 2 dpi, cellular clusters characterised by an increased cell density and stronger nucleic acid staining (Fig 3B, yellow arrows) coexisted with areas with normal nucleic acid staining levels but reduced cell-to-cell contact (Fig 3B, white arrows). By 3 dpi, a significant loss of cell confluency was observed (Fig 3C, Fig S6B-D), with large cell clusters no longer visible. To quantify these changes across multiple areas of the chip, we enumerated the number of nuclei per unit area of the membrane surface for infected LoCs at 1 and 3 dpi and uninfected controls. On the epithelial side, the density of nuclei declines progressively due to infection irrespective of the presence (p=0.010, p=9E-4) or absence (p=0.054, p=0.029) of macrophages compared to uninfected controls (Fig. 3E). In contrast, the density of cellular nuclei increases in the endothelial layer at 1 dpi before decreasing back to levels in the control samples at 3 dpi (Fig. 3F), consistent with our observation of the transient formation of endothelial cell clusters. As a further measure of quantification, we measured the proportion of the surface area of the endothelial cell layer with low or no actin staining using a range of intensity thresholds (Fig. S2E). At 3 dpi, a much larger proportion of the surface area of the endothelial layer of infected LoCs had low or no actin staining compared to an uninfected control for all intensity thresholds (Fig. S2E). Immunostaining for the viral S protein at 3 dpi (Fig 3G for endothelial and Fig 3H for epithelial layers, additional examples in Fig. S6B-D, S6F and controls with secondary antibody only in Fig. S8D-G) show clear evidence of individual infected epithelial (Fig. 3I) and endothelial cells (Fig. 3J). In some cases, S proteins appear to co-localize with the periphery of the nucleus (Fig. 3I, 3J, white arrows). The transfer of viral proteins from the apical to vascular channel and difference in confluency between the epithelial and endothelial layers at 3 dpi can be better appreciated from a 3D view of the two channels together with the connecting pores in the PDMS membrane; S protein can be clearly localised to the interface of the two cell types in the pores of the membrane (yellow arrows, Fig, 3K). Epithelial and endothelial cells exhibit very different physiological responses to low levels of infection, and vascular damage by 3 dpi was a consistent observation across all infected LoCs (n=12) maintained until 3 dpi in this study

**Fig. 3.**
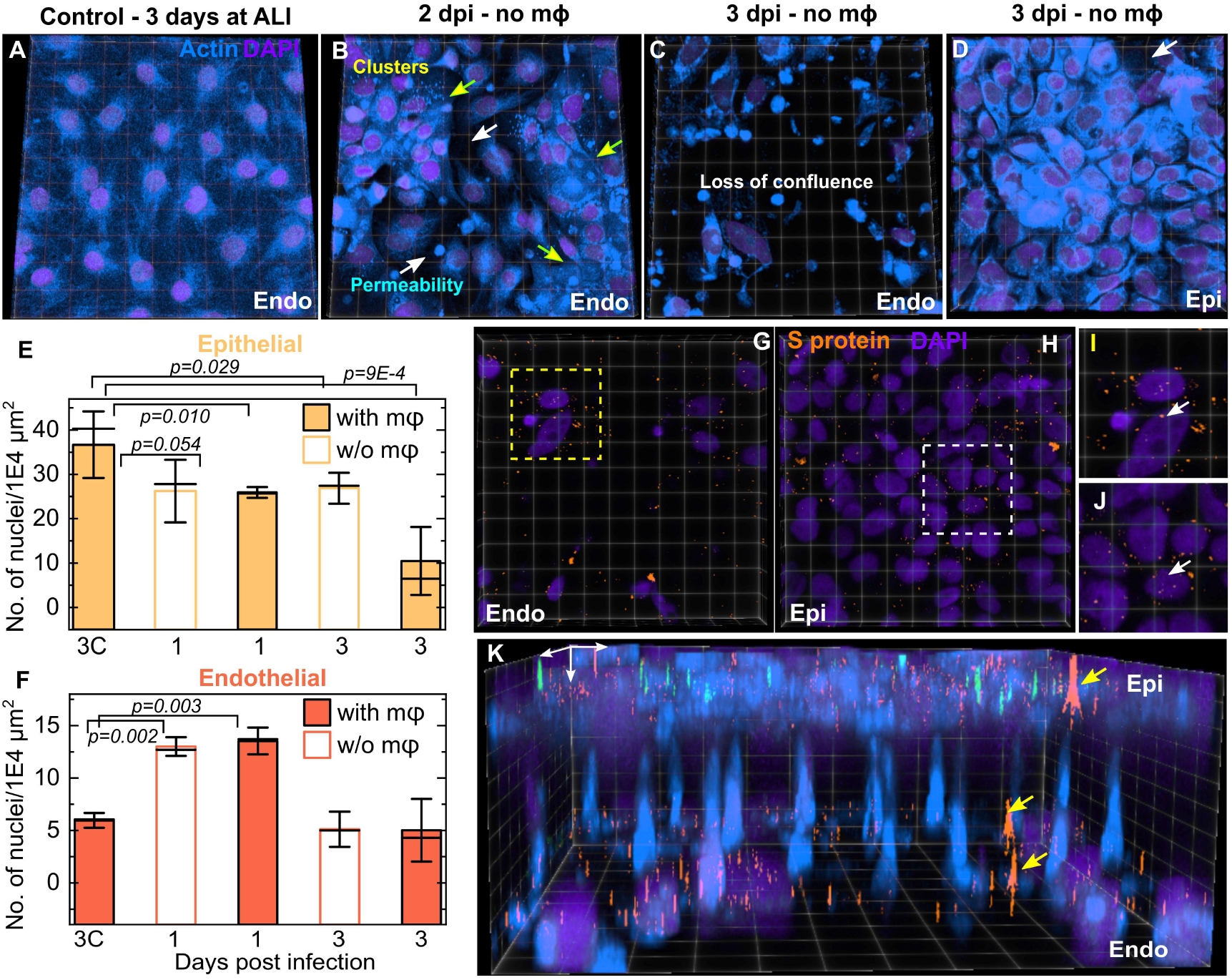
SARS-CoV-2 infection rapidly disrupts confluency of the endothelial layer. (**A-D**) 3D views of representative 232 × 232 μm^2^ (**A, B**) and 155 × 155 μm^2^ (**C, D**) fields of view from confocal imaging of an uninfected control (**A**), and infected chips without macrophages at days 2 (**B**) and 3 (**C, D**) post infection respectively. Actin and nuclear labelling are shown in azure and indigo, respectively. (**B**) At 2 dpi, adjacent areas of endothelial cell clusters identified by a stronger nuclear stain and brighter actin staining (yellow arrows) and cells with increased permeability and loss of confluence (white arrows) are indicated. (**C**) At 3 dpi, a significant loss of barrier integrity is observed in the endothelial layer, whereas the epithelial layer directly above this field of view (**D**) is relatively intact. A small area of missing cells in the epithelial layer is indicated with a white arrow in (**D**). Plots of density of nuclei (number of nuclei per 1E4 μm^2^) on the epithelial (**E**) and endothelial (**F**) faces of infected LoCs both with and without macrophages for characterization of the cellular layer integrity. The bars represent the mean value, the solid line represents the median and the error bars represent the standard deviation from at least 4 fields of view each from two independent LoCs for each timepoint. Data from uninfected controls corresponding to the 3 dpi timepoint are labelled ‘3C’. The fields of view in (**C, D**) are shown also in (**G, H**) respectively, SARS-CoV-2 spike (S) proteins labelled via immunofluorescence are false-colored in amber. Zooms in (**I**) and (**J**) show clear evidence of intracellular spike protein within both endothelial and epithelial cells, respectively. (**K**) A 3D view of the field of views in (**C, D**) and (**G, H**) respectively highlights the relative damage to the endothelial layer versus the epithelium, as well as the basolateral transmission of S protein via junctions in the pores of the membrane (identified with white arrows). P-values are calculated using a one-way Kruskal-Wallis ANOVA test.

**Fig. 4.**
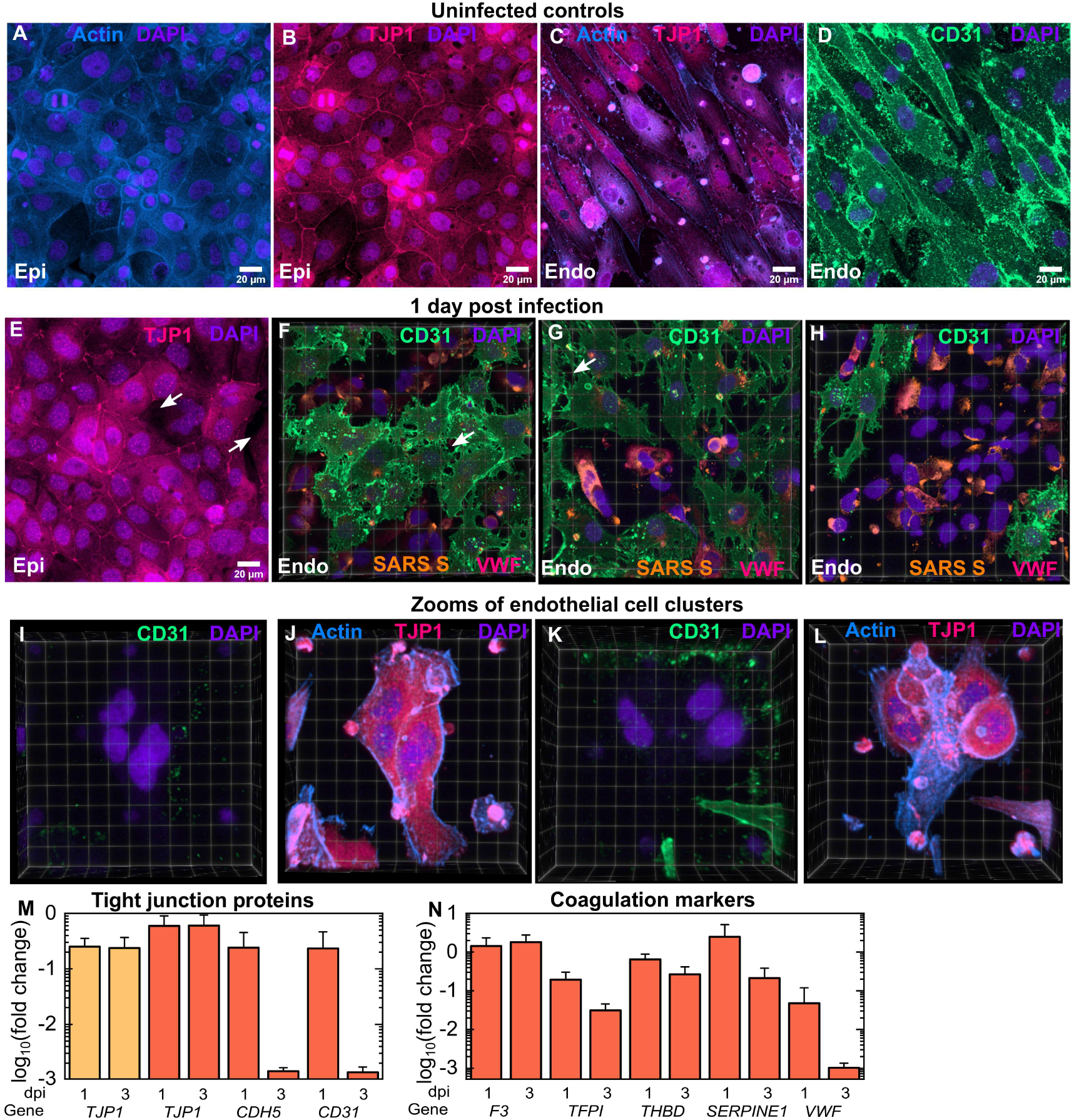
SARS-CoV-2 infection leads to a loss of tight junctions, generation of CD31-endothelial cell clusters and a pro-coagulatory environment. (**A-D**) Maximum intensity projects of 232 × 232 μm^2^ fields of view from the epithelial (**A**, **B**) and endothelial layers of an uninfected control LoC. Epithelial cells show strong actin (**A**) and TJP1 expression (**B**, identified via antibody staining) at the cell junction boundaries. Endothelial cells also show high levels of TJP-1 expression (**C**) together with CD31 expression at cell junctions (**D**, identified via antibody labelling). Actin, TJP-1, CD31, and nuclear labelling are indicated with the azure, bright pink, spring green and electric indigo LUTs respectively. 3D views of one 232 × 232 μm^2^ field of view of the epithelial (**E**) and (**F**-**H**) three 232 × 232 μm^2^ fields of view of the endothelial layers of an infected LoC at 1 dpi. Areas with lower TJP1 expression in the epithelial layer in (**E**) are indicated by white arrows. (**F-H**) Areas with loss of tight junctions between cells on the endothelial surface is indicated by white arrows. The proportion of the surface area covered by confluent cells decreases sequentially from (**F**) to (**H**). Endothelial cell clusters with no CD31 expression are clearly visible and the proportion of the surface area covered by these clusters increases sequentially from (**F**) to (**H**). In (**F**-**H**), spike protein is identified via an anti-SARS-spike antibody and shown in amber LUT, and von Willebrand Factor identified by antibody labelling is shown in bring pink LUT. Zooms of two endothelial cell clusters (**I, J** and **K, L**) in the endothelial layer of another infected LoC at 1 dpi confirm that little or no CD31 expression is detected (**I**, **K**) despite high levels of actin and TJP-1 protein expression in these cells. (**J**, **L**). (**M**) Plot of the fold change in expression of *TJP1* (epithelial and endothelial layers) and *TJP1*, *CDH5*, and *CD31* (endothelial layers only) in infected LoCs at 1 dpi (n=3) vs, uninfected controls (n=3). (**N**) Plot of the fold change in gene expression of endothelial cell specific coagulation markers - Tissue Factor (*F3*), Tissue Factor Pathway Inhibitor (*TFPI*), Thrombomodulin (*THBD*), Plasminogen Activator Inhibitor (*SERPINE1*) and von Willebrand Factor (*VWF*) in infected LoCs at 1 dpi (n=3) vs. uninfected controls (n=3). In all plots, bars represent the mean value and error bars represent the standard deviation.

We next sought to understand if these responses were specific to the lung-on-chip physiology and co-culture, and so examined infections of epithelial and endothelial cell monoculture. At 1 dpi, heavily infected epithelial cells in monoculture with 1000 PFU showed similar levels and intracellular distributions of spike protein (Fig. S5B). In stark contrast, monoculture infections of endothelial cells with viral innocula up to 100-fold greater than the apical inoculum used in the LoC experiments (Fig. 2A) showed few foci of viral S proteins. Endothelial cells retained their morphology and confluency (Fig. S7) up to 2 dpi, in agreement with recent reports of the inability of apical inoculations of SARS-CoV-2 to infect lung microvascular endothelial cells ^18^. This suggests that the site of contact of SARS-CoV-2 with endothelial cells (apical vs basolateral) may play an important role – a physiological factor that can be recapitulated in the LoC but not in simpler models.

### SARS-CoV-2 infected endothelial cells lose expression of tight junction markers and adopt a pro-coagulatory phenotype

Actin staining definitively demonstrated the vascular damage caused as a result of loss of cell confluence, we therefore probed cells from both layers in infected and uninfected LoCs for changes in expression of tight junction proteins that might have caused these changes Maximum intensity projections from the epithelial cell layer of an uninfected control showed strong expression of both actin (Fig. 4A) and TJP1 at the cell junctions (Fig. 4B). Similarly, images from a confluent endothelial cell layer in an uninfected control LoC showed actin staining correlated with high expression of Zonula Occludens − 1 (ZO-1 or TJP1) (Fig. 4C) as well as high expression of the endothelial cell specific marker CD31 at cell junctions (Fig. 4D). These results were corroborated by qRT-PCR measurement of high levels of expression of *TJP1* and *CD31* as well as that of VE-cadherin (*CDH5*) in uninfected controls (Fig. S8A).

Consistent with Fig. 3, changes to the expression pattern of TJP1 in the epithelial layer at 1 dpi were subtle, TJP1 expression at cell junctions continued to be observed across much of the epithelial cell layer (Fig. 4E, additional images in Fig. S8B, C). SARS-CoV-2 infection, on the other hand, profoundly altered CD31 expression and distribution in the endothelial layer. 3D views of regions of interest in the endothelial layer of an infected LoC showed this to be a dynamic process that resulted in two significant changes: first, an increase in vascular permeability from the loss of tight junctions between CD31+ cells (white arrows in Fig. 4F, G), and second, the presence of cells that showed little or no CD31 expression (Fig. 4F-H). Cells displaying the ‘CD31-phenotype’ typically had higher levels of spike protein (shown in amber in Fig. 4F-G), and an increased local cell density and nuclear labelling consistent with the endothelial cell clusters identified in Figure 3. This link is strengthened by examination of zooms of two endothelial cell clusters (Fig. 4I, J, and Fig. 4K, L) from another LoC at 1 dpi. This confirmed that cells in both clusters were CD31- (Fig. 4I, Fig. 4K) but nevertheless express TJP1 and showed strong and altered actin distributions (Fig. 4J, Fig. 4L), morphological features consistent with the clusters identified in independent experiments in Fig. 3. These changes are further corroborated by qRT-PCR measurements which show a progressive and sharp reduction in overall levels of *CD31* and *CDH5* expression in the endothelial layer at 1 dpi and 3 dpi respectively compared to uninfected controls (Fig. 4M). In contrast, the reduction in *TJP1* expression at 1 dpi is less pronounced and expression levels do not decline further at 3 dpi.

These dramatic changes in expression of tight junction proteins also resulted in series of complex alterations to the expression of endothelial cell proteins with direct roles in coagulatory pathways (Fig. 4N). For example, infection did not alter expression of *pro-coagulatory* Tissue Factor (*F3*) released by endothelial cells as part of the extrinsic coagulation pathway^34^, but induced a progressive decrease in the expression of *anti-coagulatory* Tissue Factor Pathway Inhibitor (*TFPI*) ^35^. A more modest but similarly consistent decrease was also observed for the expression of Thrombomodulin (*THBD*) that also has an anti-coagulatory function. Although infection also decreased the expression of von Willebrand Factor (*VWF*), elevated levels of VWF protein were observed on the surface of the CD31-endothelial cells at 1 dpi via immunofluorescence (bright pink, Fig. 4F-G). In contrast, expression of plasminogen inhibitor activation (PAI-1 expressed by *SERPINE1*) decreased only by 3 dpi. Taken together with the loss of CD31 expression, these results point to the generation of a pro-coagulatory environment in the vascular channel of infected chips.

### SARS-COV-2 persists in individual epithelial and endothelial cells in the alveolar space

Our results thus far showed that infection on-chip itself lead to heterogenous outcomes that were strongly cell-type specific. We therefore performed a series of RNAscope assays with confocal microscopy to probe the spatial distribution and cell-to-cell variability in gene expression at the single-cell level to complement observations from qRT-PCR measurements. We began by examining levels of viral RNA and antisense RNA generated during viral replication *in situ* using probes for *S* RNA and *ORF1AB* antisense RNA respectively. Representative images for 232 × 232 μm^2^ fields of view from the apical and vascular channels of chips at 1 and 3 dpi are shown in Fig 5A-H. At 1 dpi, a few examples of heavily infected cells are visible on the epithelial layer (Fig. S9A-C) and viral genomic RNA can be detected throughout the cytoplasm. Consistent with the observations in Fig. S5, such cells are likely to be the source of infectious virions released in the apical wash in Fig. 2A, B. However, in the vast majority of cells on the epithelial layer, the virus is detected only at low levels at 1 dpi (Fig. 5A), and infection does not appear to be limited by (Fig. 5C, white arrows) or even correlated to cellular *ACE2* expression at the time of fixation (Fig. 5D, yellow arrows). Both genomic and antisense RNA are detected in endothelial cells at 1 dpi (Fig. 5B), indicating that intracellular viral replication can also occur in these cells. Heavily infected endothelial cells were not observed, in agreement with the lack of infectious virions in the vascular effluent (Fig. 2C). Endothelial cell clusters are clearly identifiable by their morphological features, and zoomed-in images (Fig 5E, F) show that viral infection of endothelial cells with normal and clustered morphology. As with the epithelial layer, instances of localization of viral RNA with the nucleus is also observed (Fig. 5F, white arrowheads). At 3 dpi, infection in both layers (Fig 5G, 5H) and the loss of endothelial layer integrity is evident (compare Fig. 5G vs. Fig. 5H). We did not detect heavily infected epithelial cells at 3 dpi, in agreement with the diminishing viral titre (Fig. 2A). We normalized the number of RNA dots detected either on a per-cell basis (Fig. 5I-K) or per-field-of-view basis (Fig. S10), to enable quantification of differences between the two layers while accounting for differences in cell numbers and cell death. The median values of intensities were similar across all conditions (Fig. S11). In the epithelial layer, viral infection and replication per cell were either similar (Fig. 5J, L) or lower by 3 dpi (Fig. 5I, p=0.018; Fig. 5K, p=0.006). In contrast, viral RNA accumulates in individual endothelial cells (Fig. 5K, Fig. 5L, p=0.004). The presence of macrophages generally leads to lower levels of viral RNA replication (Fig. 5I, J) but does not prevent the accumulation of viral genomes intracellularly in endothelial cells (Fig. 5L). At 3 dpi, *ACE2* levels are higher on a per-cell basis in endothelial vs epithelial cells (Fig. 5M, p=0.028), due to a reduction in *ACE2* levels on the epithelial layer.

**Fig. 5.**
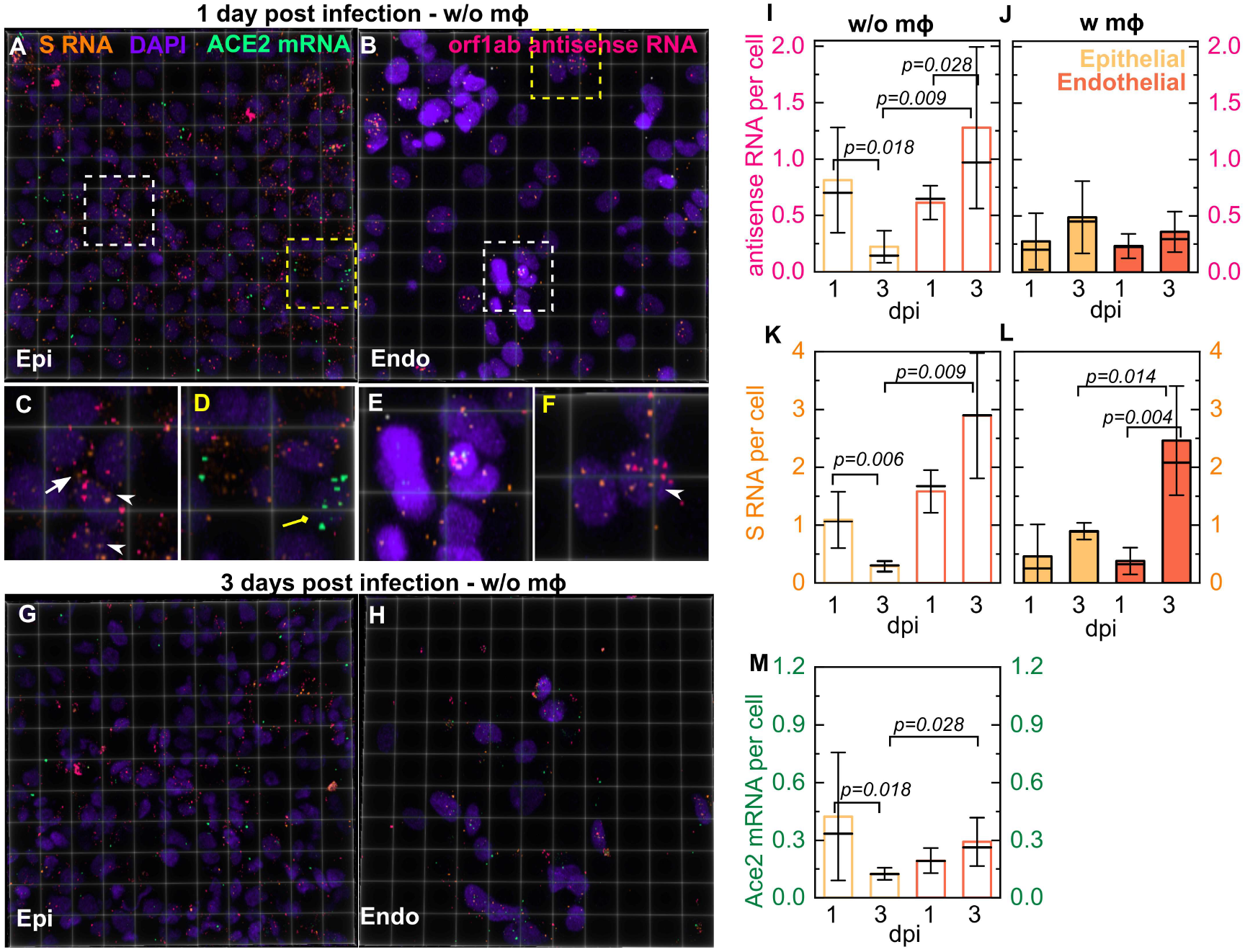
RNAscope analysis reveals intracellular localization and slow accumulation of intracellular viral RNA in the LoC model. 3D views of 232 × 232 μm^2^ fields of view of the epithelial (**A, G**) and endothelial layer (**B, H**) from infected LoCs reconstituted without macrophages at 1 (**A, B**) and 3 dpi (**G, H**). *Orf1ab* antisense RNA, *S* RNA, *ACE2* mRNA, and nuclear staining with DAPI are false-colored pink, amber, spring green, and electric indigo, respectively. (**C, D**) Zooms corresponding to the region in (**A**) highlighted with white and yellow boxes, respectively. (**C**) An example of infection and intracellular replication in a cell with no detectable *ACE2* expression (white arrow) as well as nuclear localization of viral RNA (white arrowhead). (**D**) An example of an uninfected cell with *ACE2* mRNA expression (yellow arrow). (**E, F**) Zooms corresponding to the regions in (**B**) representing increased nucleic acid staining (hyperplasic) and normal levels of nucleic acid staining highlighted with white and yellow boxes, respectively. (**E**) Examples of infection of hyperplasic endothelial cells both with and without *ACE2* expression (**F**) Examples of endothelial cell infection with no *ACE2* expression, nuclear localization of viral RNA in an infected endothelial cell is indicated (white arrowheads). Quantification of viral antisense RNA (**I, J**) and viral genomic RNA (**K, L**) from pairs of otherwise identical LoCs reconstituted without (**I, K**) and with macrophages (**J, L**) analyzed at 1 and 3 dpi. Plots show the number of spots normalized by the total number of cells in 4-6 field of views detected using RNAscope and confocal imaging using identical imaging conditions for all chips. Bars represent the mean value, the solid line represents the median, and error bars represent the standard deviation. (**M**) Quantification of *ACE2* expression per cell in epithelial cells and endothelial cells in LoCs reconstituted without macrophages at 1 and 3 dpi respectively. P-values are calculated using a Kruskal-Wallis One-Way ANOVA Test.

Normalization on a per-field-of-view basis shows predominantly a *declining* trend for levels of replication (Fig. S10A, B) and intracellular viral RNA (Fig. S10C, D) in both cell types, as well as a *decline* in *ACE2* expression (Fig. S10E), consistent with qRT-PCR measurements (Fig. 1J, Fig. S10F). These results highlight the cumulative effect of differential cell numbers and cell death in these two layers (Fig. 3E, F). Overall, virus levels are low and decline in both layers by 3 dpi, yet the virus persists and continues to replicate within individual infected cells, in agreement with reports of clinical infection.

### Persistent NF-KB inflammatory responses to SARS-CoV-2 infection in endothelial but not epithelial cells

Concurrently, we investigated host responses to the infection. Expression levels of NF-KB related pro-inflammatory genes (*TNFA*, *IL6*, *IL1B*) at 1 dpi was an order of magnitude higher than that of interferon genes in both cell types, with little to no interferon stimulatory gene expression detected in both cell types (Fig. 6A). Responses were also cell-type specific; endothelial cells from infected LoCs showed upregulation of *TNFA*, *IL6* and *IFNB*, *IFNL1* and *IFNL3* genes whereas only expression of *IFNB* and *IFNL1* in epithelial cells were higher than uninfected controls (Fig. 6B). In both cell types, expression of *IP10* and *ISG15* were lower than uninfected controls, consistent with the action of a number of SARS-COV-2 proteins in inhibiting type I interferon signalling ^36,37^.

**Fig. 6.**
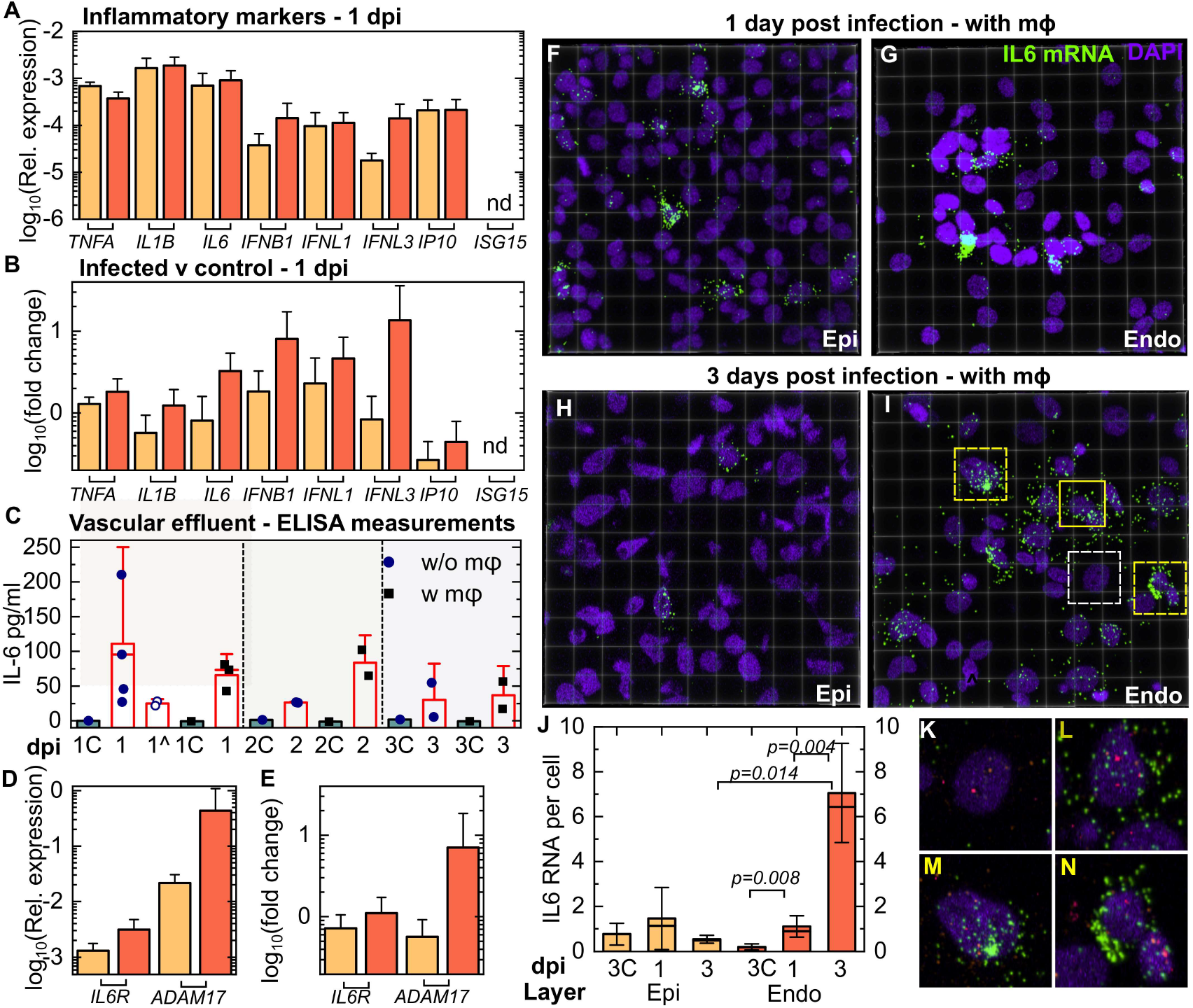
SARS-CoV-2 infection generates a persistent pro-inflammatory response in endothelial cells. (**A**) Expression relative to *GAPDH* of pro-inflammatory cytokines (*TNFA, IL1B, IL6)*, interferon genes (*IFNB, IFNL1, IFNL3*) and interferon stimulating genes (*IP10, ISG15*) in the epithelial and endothelial layer of infected LoCs at 1 dpi reconstituted without macrophages (n=3 for *IFNL1* and *IFNL3* in the epithelial layer, n=4 for all other data). (**B**)Fold-change in the markers in (**A**) relative to uninfected controls (n=3) (data in Fig. S12) at the same timepoint. The bar represents the mean and the error bars the standard deviation. (**C**) Enzyme-linked immunosorbent assay (ELISA) measurements for IL-6 in the vascular effluent for at least two infected LoCs at each timepoint reconstituted with and without macrophages. Uninfected controls for each condition are labelled with a ‘C’ and grey bars, and LoCs with Tocilizumab administration are indicated by ^. The height of the bars represents the mean, the median is represented by a solid line, and the error bars represent the standard deviation. Each dot represents the mean of two technical replicates. (**D**) Expression of *IL6R* and *ADAM17* relative to *GAPDH* in the epithelial and endothelial layer of infected LoCs at 1 dpi (n=3 for the epithelial layer and n=4 for endothelial layer). (****E****) Fold-change in expression relative to uninfected controls (n=3 for both layers). (**F-I**) 3D views of representative 232 × 232 μm^2^ fields of view of the epithelial (**F, H**) and endothelial layer (**G, I**) from infected LoCs reconstituted with macrophages at 1 (**F, G**) and 3 dpi (**H, I**). *IL6* mRNA, and nuclear staining are indicated with the chartreuse and electric indigo LUTs respectively. (**K-N**) Zooms corresponding to the regions in (**I**) highlighted with white (**K**) and yellow boxes with solid (**L**) and dashed lines (**M**, **N**) respectively. In these panels, *orf1abantisense* RNA (pink) and *S* RNA (amber) are also shown. The panels show examples of cells with similar levels of viral infection but with no (**K**), intermediate (**L**) and high levels (**M, N**) of *IL6* expression. (**J**) Quantification of *IL6* expression in epithelial and endothelial cells from a pair of otherwise identical LoCs analyzed at 1 and 3 dpi, respectively. Plots show the total number of spots normalized by the number of cells in 4-6 fields of view detected using RNAscope assay using identical imaging conditions for all chips. Bars represent the mean value, the solid line represents the median, and error bars represent the standard deviation. Data from uninfected controls for 3 days at air-liquid interface is indicated by ‘C’. The bar represents the mean and the error bars the standard deviation. P-values are calculated using a Kruskal-Wallis One-Way ANOVA Test.

The LoC platform also allows the simultaneous detection of secreted cytokines in the vascular effluent from the same chips. ELISA assays for IL-1B and IP10 did not detect these cytokines in the effluent (data not shown), the lack of IL-1B secretion despite high expression (Fig. 6B), suggests that it is not exported^38^. IL-6, on the other hand, was consistently detected up to 3 dpi in the vascular effluent from LoCs both in the presence or absence of macrophages (Fig. 6C). No IL-6 was detected in the vascular effluent from uninfected controls. IL-6 levels were not significantly different between these two categories of LoCs, which strongly suggested a non-immune cell source for this cytokine (Fig. 6C).

Given the pleiotropic nature of IL-6^39^, we also examined expression of the IL6R receptor and the metallopeptidase ADAM17 which sheds the TNF-alpha receptor and the IL-6R receptor^40^ in both the epithelial and endothelial layer at 1 dpi (Fig. 6D). Expression of *ADAM17*, which has been shown to enhance vascular permeability^41^, was on average 7-fold higher in the endothelial layer at 1 dpi (Fig. 6E), suggestive of the activation of the IL-6 signalling pathway.

COVID-19 patients with severe infection have elevated IL-6 levels, and we therefore sought to examine the cell-type specific *IL6* expression at the single cell resolution. We therefore used RNAscope to probe the spatial distribution of *IL6* expression in LoCs reconstituted with macrophages. Consistent with the qRT-PCR data (Fig. S12), *IL6* expression in the control LoC is higher in the epithelial layer (Fig. S13A) than the endothelial layer (Fig. S13B), and immunostaining with an anti-CD45 antibody confirmed that the *IL6* expression in the epithelial layer of the chip did not co-localise with macrophages (Fig. S13A, B). At 1 dpi, individual cells with higher levels of *IL6* expression were observed in both epithelial and endothelial layers (Fig. 6F, G), but only differences in expression in the endothelia layer were statistically significant (Fig. 6J). Cells with high *IL6* expression were also unlikely to be macrophages (Fig. S13 C-E). By 3 dpi *IL6* expression in the epithelial layer had returned to the control levels (Fig. 6H, J, S14A) whereas expression increased dramatically and was more widespread in the endothelial layer (Fig. 6I, J, S14A, S14B). Expression did not appear to correlate with level of infection – zooms of cells with comparable levels of infection in Fig. 6I with low (Fig. 6K), medium (Fig. 6L) and high (Fig. 6 M, N) *IL6* expression are shown.

Expression in the endothelial layer at 3 dpi was higher than that in the endothelial layer at 1 dpi, the epithelial layer at 3 dpi, and the uninfected controls, both on a per-cell and per-field-of-view basis (Fig. 6J, Fig. S14A, p=0.004, p=0.014, p=0.008 respectively). Unlike viral RNA levels (Fig. S10B, D) *IL6* expression in the endothelial layer *does not diminish* over 1-3 dpi and would therefore appear to be the major contributor to IL-6 secretion in the vascular effluent as infection progresses.

### Inhibition of trans IL-6 signalling reduces but does not eliminate endothelial cell inflammation

The activation of IL-6 signalling observed suggested *trans* IL-6 signalling as a potential target to ameliorate the vascular inflammation observed. A similar rationale has led to numerous clinical trials of the anti-IL-6R monoclonal antibody Tocilizumab^42^ as a repurposed therapeutic for COVID-19. Tocilizumab administration at 56 μg/mL via continuous perfusion over 2 days post infection *per se* did not abrogate IL-6 secretion, consistent with its mode of action (Fig. 6C). We therefore undertook an investigation of cell morphologies in the endothelial layer of a treated and untreated LoC via actin staining. This revealed that whereas endothelial cell confluency was, on the whole, better retained in the treated chip (Fig. 7A, zooms in Fig. 7B, untreated control in Fig. 7D, zoom in Fig. 7E) Tocilizumab administration *did not prevent* the occurrence of endothelial cell clusters (Fig. 7A, zooms in Fig. 7C, untreated control in Fig. 7D, zoom in Fig. 7F). Consistent with these results, immunostaining for CD31 in another treated LoC showed an improvement in the integrity of the endothelial cell layer with tight junctions observable at 2 dpi (3D views in Fig. 7G, H) yet endothelial cell clusters with reduced CD31 expression and strong actin staining were nevertheless formed (3D views in Fig. 7G, H, indicated by yellow arrows).

**Fig. 7.**
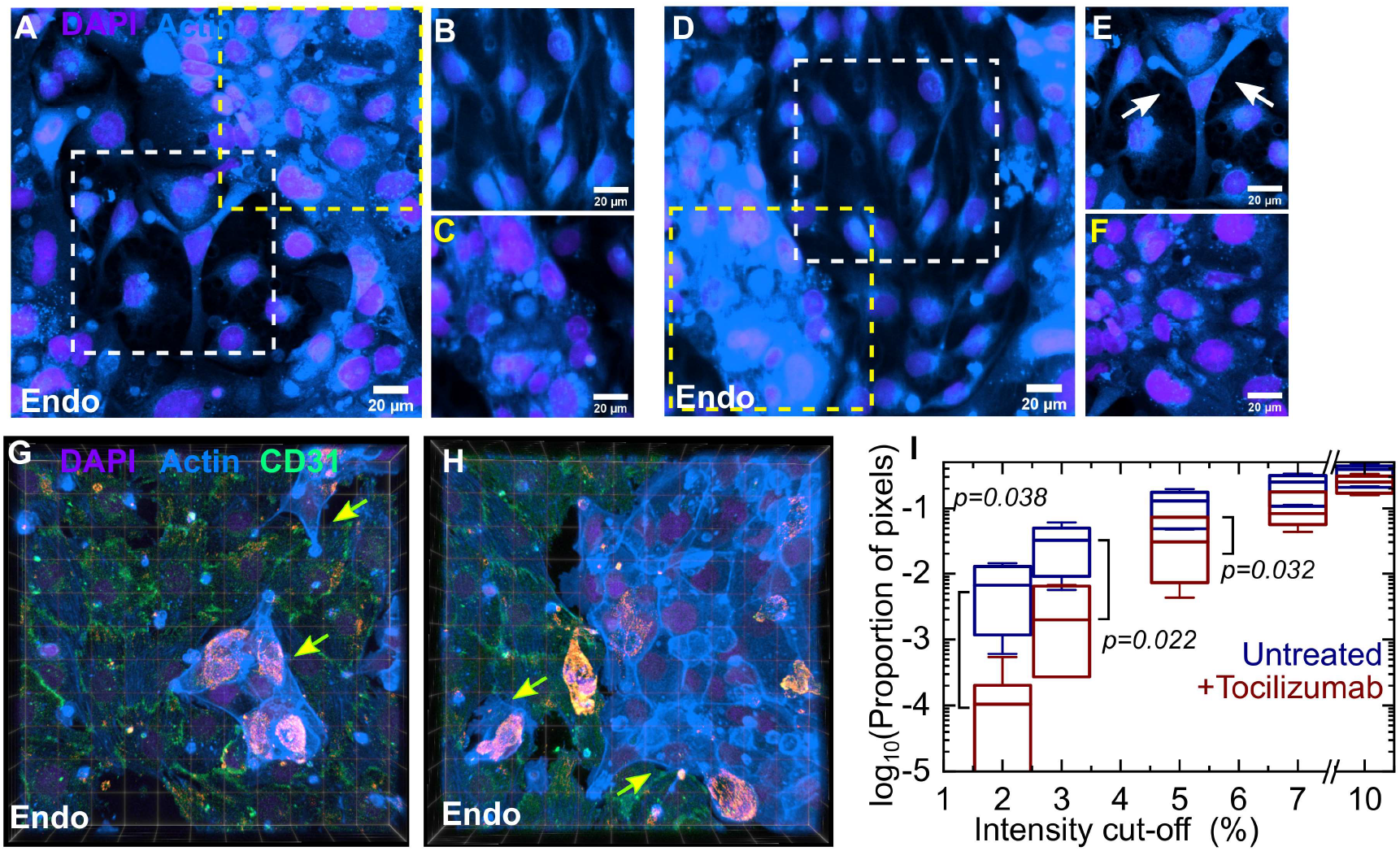
Tocilizumab treatment reduces vascular permeability but does not prevent formation of endothelial cell clusters with reduced *CD31* expression. Maximum intensity projections of representative 232 × 232 μm^2^ fields of view of the endothelial layer from an infected LoC reconstituted without macrophages at 2 dpi with (**A**) and without Tocilizumab treatment (**D**). Actin and nuclear staining are indicated with the azure and electric indigo LUTs respectively, the actin staining is saturated for the endothelial cell clusters to highlight cells with lower levels of actin. (**B, C**) Zooms corresponding to the region in (**A**) highlighted with white and yellow boxes, respectively. (**B**) A region with normal vascular profile, (**C**) A region of endothelial cell cluster formation with increased actin staining. (**E, F**) Zooms corresponding to the regions in (**D**) representing regions highlighted with white and yellow boxes, respectively. (**E**) An example of reduced tight junctions between endothelial cells; bare patches are indicated with arrows. (**F**) A region of endothelial cell clusters. (**G**) 3D views from two 232 × 232 mm2 field of views of the endothelial layer of a Tocilizumab treated infected LoC at 2 dpi, immunostaining for the tight junction protein CD31 is shown in the Spring Green LUT. Endothelial cell clusters with low CD31 expression and altered actin staining are also visible and indicated by yellow arrows. (**I**) A plot of the proportion of pixels (shown on a logarithmic scale) with regions of low F-actin intensity identified via cut-off thresholds defined as a percentage of the maximum intensity in 6-7 regions of interest each for LoCs with and without Tocilizumab administration. ROIs were chosen from 232 × 232 μm^2^ fields of view and were defined to exclude the occurrence of hyperplasic cells. A significantly larger fraction of the surface area of the ROIs for the untreated chips have low F-actin intensities. P-values are calculated using a Kruskal-Wallis One-Way ANOVA Test.

To quantify the improvement in permeability, we compared regions of interest (ROIs) that excluded areas of cellular hyperplasia across at least six fields of view from the endothelial layer of LoCs with and without Tocilizumab perfusion, and identified areas with low or no F-actin staining as those with reduced confluence, as in Fig. S6E. A plot of the proportion of pixels with intensities below a defined cut-off threshold (Fig. 7I) showed that the untreated LoC did have a significantly higher proportion of pixels with intensities lower than 5% (p=0.032) of the maximum intensity (p=0.022 for 3% and p=0.038 for 2%). Inhibition of IL-6 signalling through Tocilizumab is therefore able to ameliorate *some* but *not all* of the vascular damage observed.

## Discussion

The alveolar space has a strikingly different physiology from that of the upper airway. ACE2 is the canonical entry receptor used by SARS-CoV-2, yet is scarce in the alveoli. It has been speculated that high levels of *ACE2* expression may be necessary for export of mature virions ^43^ and because most virology assays measure viral titre, studies with alveolar epithelial cell lines such as A549 have focused on increasing *ACE2* expression through transfection. In contrast, alternate receptors such as NRP1 more abundant in the lower airways have been understudied, and the effects of uptake via NRP1 vs. ACE2 on the viral life cycle is relatively unknown. Here, we populate our model with relevant primary human cells and report low *ACE2* expression and high *NRP1* expression, consistent with human alveolar physiology. Similarly, most infections of alveolar epithelial cells are usually performed at high MOI^16^. Instead, we used an infectious dose that better mimics either a direct inspiration of aerosols or the transfer of infectious virions from the upper respiratory tract. These features, together with the ability to co-culture epithelial cells (at an air-liquid interface) and endothelial cells (under flow) enable the lung-on-chip model to be used to study aspects of early infection that cannot be achieved with other systems.

Despite low ACE2 expression and low MOI, SARS-CoV-2 infection produced pronounced physiological changes on-chip. This could be due to the high levels of NRP1 and aided by upregulation of *TMPRSS2*, which is known to be important in human SARS-CoV-2 infection. In contrast to models of productive replication, we observe slow intracellular viral replication without significant release of infectious virions, observations that are consistent with reports of recovery of viral RNA in patients after they cease to be infectious^44^. Microscopy-based analyses also revealed that responses to infection are highly heterogenous; for example, the few foci of heavily infected cells might represent cells with constitutively higher levels of *ACE2* expression at the time of infection. However, why alveolar epithelial cells show persistent but not productive infection is still unclear.

Infection of the epithelial layer rapidly led to an infection of the endothelial layer. Basolateral transmission cannot occur in typical monoculture experiments, but it has not been reported to occur to a large extent in monocultures of airway epithelial cells at the air-liquid interface. However, the basement membrane is much thinner in alveoli than in the upper airway^45^, with cell-to-cell contact between the epithelial and endothelial cell layers that is recreated in the array of pores on-chip. Infection of lung microvascular cells therefore might occur through uptake of vesicles as part of cell-to-cell communication between the layers (a receptor-independent mechanism of uptake). These observations are strengthened by the lack of observation of any signs of infection in lung microvascular endothelial cells when directly exposed to the virus on their apical surface. Alternatively, infection of endothelial cells when exposed to the virus basolaterally might still occur due to receptor-mediated uptake, with differences between apical and basolateral infection explained by differences in distribution of cell surface receptors. Notably, NRP1 is likely to be located along the basal surface of the endothelial cells as an integrin-binding protein^26^, and SARS-CoV-2 has been shown to infect vascularised organoids, albeit at high MOI^46^. The exact mechanisms remain to be elucidated, but the consequences of this may extend downstream to the persistent infection and endothelial-cell specific cell damage observed.

A striking feature of SARS-CoV-2 infections in the LoC model was the cell-type specific morphological changes. Infection altered endothelial cell morphology in two different but interconnected ways. First, the loss of tight junctions between cells is indicative of increased vascular permeability that could lead to exudative alveolar damage observed in autopsy reports. Exposed sub-endothelial collagen would likely trigger the contact activation or intrinsic pathway of coagulation. This was also accompanied by pro-coagulatory changes in the extrinsic pathway through a reduction in expression of anti-coagulatory factors such as TFPI and thrombomodulin. Interestingly, infection of Human Umbilical Vein Endothelial Cells (HUVECs) with Dengue virus has been shown to have the opposite effect ^47^, with a strong increase in thrombomodulin expression consistent with the disease manifestation as a haemorrhagic fever. These results highlight the close link between coagulation and inflammation in this cell type. Second, a subpopulation of endothelial cells completely lost CD31 expression and adopted an altered morphology consisting of cell clusters. These clusters could form either through enhanced proliferation or from cell migration.

The former is more likely, as a consistent increase in cell density was observed across multiple LoCs, but both possibilities are consistent with the role of NRP1 as a growth factor receptor or its role in VEGF signalling^48^. The loss of CD31 expression could itself have a number of possible implications: CD31 itself plays important roles in leukocyte migration^49^, angiogenesis, in maintaining a anti-coagulatory environment^50,51^ as well as preventing endothelial cell death from immune cell mediated inflammation^52^. These CD31-cells could further contribute to a pro-coagulatory environment in the microvasculature and lead to changes in vascular architecture reported in autopsies^11^. A loss of CD31 expression in endothelial cells will likely alter interactions with neutrophils and platelets that who also express CD31, potentially contributing to the formation of neutrophil extracellular traps and platelet hyperactivity reported in COVID-19^53,54^. These findings are also consistent with the observation of elevated serum levels of CD31 in patients with severe COVID-19^55^. The exact mechanisms for the loss of CD31 and other tight junction markers remains to be elucidated, but a possible mechanism could be through action of viral proteins themselves, as is observed in the case of the K5 protein of the Kaposi Sarcoma herpesvirus^56^ that causes Kaposi Sarcoma.

Increased NF-KB inflammatory responses could also account for the vascular changes observed, and the role of cytokine storms in COVID-19 pathogenesis is still unclear. We observed that low levels of intracellular viral RNA generates an NF-KB mediated pro-inflammatory response, with suppression of antiviral interferon responses, consistent with reports of the effect of a number of viral proteins^57,58^. Within each cell types, there were significant cell-to-cell differences in expression of inflammatory markers, clearly identifiable with single-cell spatial resolution through the use of RNAscope and confocal imaging in the LoC model. These differences could be due to differences in the mode of uptake, or differences in the intracellular localization of viral RNA and RNA replication, or differences in the levels of translation of viral proteins. We observe instances of co-localisation of viral RNA and S protein with the nucleus, which is consistent with bioinformatic predictions using algorithms for RNA localisation^59^, but has not been reported in studies using cell lines with productive infections. These atypical sub-cellular localisations may be characteristic of persistent infection and merit further investigation. Yet, once again the responses of epithelial and endothelial cells contrasted sharply – inflammation in epithelial cells was transient, whereas it was persistent in endothelial cells.

Lastly, given the persistent *IL6* expression in endothelial cells, we applied the model and our microscopy-based approach to examine if Tocilizumab perfusion could the occurrence of either increased vascular permeability or the formation of endothelial cell clusters or both. We found that Tocilizumab *was* able to maintain cellular confluency but did not prevent a subset of endothelial cells from over-proliferating and acquiring an inflammatory phenotype. These observations at the single cell level offer insights into reasons why outcomes with this drug in clinical trials have been modest and at times contradictory^60^ and there is insufficient evidence of the efficacy of Tocilizumab in reducing mortality. The lung-on-chip model is therefore able to make clinically testable predictions, highlighting the utility of this approach.

In sum, SARS-CoV-2 infection of a model of the alveolar space shows a set of unique characteristics underlined by persistent infection, low viral replication, inflammatory response, damage to endothelial barrier integrity, and the formation of a pro-coagulatory environment that are in good agreement with reports of clinical disease. The superior physiological mimicry LoC model (air-liquid interface, co-culture, perfusion through a vascular channel) and high-resolution confocal imaging reveals insights into the dynamics and mechanisms that underlie these observations that are difficult to obtain in other experimental models and that will improve our understanding of the pathogenesis of this multi-organ disease.

## Acknowledgements

We gratefully acknowledge assistance from Dr Muhammet Fatih Gulen for the ELISA assays. We are also grateful to the members of the BioImaging Core Facility (BIOP) for assistance with confocal microscopy and to Prof Dr Thomas Hugle at the Centre Hospitalier Universitaire Vaudois (CHUV(for donation of Tocilizumab.

## Methods

### Cell culture

Primary human alveolar epithelial cells (epithelial cells) and human lung microvascular endothelial cells (endothelial cells) were obtained from a commercial supplier (Cell Biologics, USA). Each vial of epithelial cells comprised a mix of Type I and Type II cells at passage 3, and expression of cell-type specific markers (*AQP5, PDPN,* and *CAV1* for type I and *ABCA3* and *SFTPC* for type II) was verified via qRT-PCR on-chip (Fig. S3D). Both cell types were cultured *in vitro* in complete medium comprising base medium and supplements (Cell Biologics, USA) in 5% CO2 at 37 °C. Passage of epithelial cells lowered *ACE2* expression (Fig. S2A), therefore all chips were reconstituted with epithelial cells seeded directly on the lung-on-chip (see below), without any additional *in vitro* culture. In contrast, we observed no significant changes in viral entry factor expression in endothelial cells upon passage, and so therefore these cells were passaged between 3-5 times before seeding in the LoC devices.

### PBMC isolation and macrophage differentiation

Peripheral blood mononuclear cells were obtained from buffy coat (Interregional Blood Transfusion SRC Ltd, Switzerland) obtained from anonymised donors. PBMCs were isolated using a Biocol Separation procedure as per the manufacturer’s instructions. Isolated PBMCs were subsequently cryopreserved in a solution of 70% heat inactivated Fetal Bovine Serum (FBS, Gibco), 20% RPMI Medium (Gibco) and 10% dimethlysulfoxide (DMSO). One week prior to seeding the macrophages in the LoC devices, a cryopreserved aliquot was thawed and cultured in a T-75 flask (TPP, Switzerland) in RPMI supplemented with 10% FBS. CD14+ monocytes were isolated using positive selection (CD14 ultrapure isolation kit, Miltenyi Biosciences) and cultured in RPMI medium supplemented with 10% FBS and differentiated for 7 days with 20 ng/mL recombinant human Macrophage-Colony Stimulating Factor protein (M-CSF) (Thermo Fisher Scientific), and 100U/L of penicillin-streptomycin solution (Thermo Fisher Scientific) to avoid bacterial contamination. The cells were cultured in plastic petri dishes without pre-sterilisation (Greiner Bio-One) so that differentiated macrophages could be more easily detached.

### Quantitative Real-Time PCR (qRT-PCR) for cell characterization

Epithelial cells obtained directly from the supplier were grown overnight in cell-culture microdishes (Ibidi) or T-25 cell culture flask (TPP, Switzerland). Passaged epithelial cells were grown to confluency in a T-75 cell culture flask (TPP). Growth media was removed from the flask, and the cells were incubated with the appropriate volume of RNA lysis buffer (RNAeasy Plus Micro Kit, Qiagen) and RNA isolated as per the manufacturer’s instructions and resuspended in 25 μL of DEPC-treated water. Approximately 1 μg of RNA was then used to generate cDNA using the SuperScript®IV First-Strand Synthesis System with random hexamers (Invitrogen), which was stored at −20 °C. Specific primers for used for the qRT-PCR are listed in Table S1. qRT-PCR reactions were prepared with SYBR®Green PCR Master Mix (Applied Biosystems) with 500 nM primers, and 1 μL cDNA. Reactions were run as absolute quantification on ABI PRISM®7900HT Sequence Detection System (Applied Biosystems). Amplicon specificity was confirmed by melting-curve analysis.

### Quantitative Real-Time PCR (qRT-PCR) for quantification of viral genomes in the supernatant

150 μL of cell-free supernatant was processed using a kit to extract viral RNA as per the manufacturer’s instructions (NucleoSpin Dx Virus, Machery Nagel). For the vascular effluent, this was taken from the approximately 1.5 mL of effluent generated daily by the flow through the vascular channel. In the case of the apical wash, the 30μL collected from the apical channel was further diluted in cell culture media up to 150μL and subsequently processed. Viral RNA was eluted in 50mL of DEPC-treated water. 5 mL of RNA was then used in a one-step qRT-PCR kit for SARS-CoV-2 detection (2019-nCoV TaqMan RT-PCR kit, Norgen Biosciences) using primer and probe mixes corresponding to the nucleocapsid (N2) gene and the host RNAseP genes. Primer sequences are those listed by the US Centre for Disease Control (CDC).

### Generation of SARS-CoV-2 stocks

VeroE6 cells and a clinical isolate of SARS-CoV-2 were a kind gift from the lab of Prof Carolyn Tapparel at the University of Geneva. SARS-CoV2/Switzerland/GE9586/2020 was isolated from a clinical specimen in the University Hospital in Geneva in Vero-E6. Cells were infected and supernatant was collected 3 days post infection, clarified, aliquoted and frozen at −80 °C and subsequently titrated by plaque assay in Vero-E6. Virus used for all experiments in this manuscript was at passage 3 in Vero E6 cells.

### Human lung-on-chip (LoC) model

LoC made of polydimethylsiloxane (PDMS) were obtained from Emulate (Boston, USA). Extracellular matrix (ECM) coating was performed as per the manufacturer’s instructions. Chips were activated using ER-1 solution (Emulate) dissolved in ER-2 solution at 0.5 mg/mL (Emulate) and exposed for 20 minutes under UV light. The chip was then rinsed with coating solution and exposed again to UV light for a further 20 minutes. Chips were then washed thoroughly with PBS before incubating with an ECM solution of 150 μg/mL bovine collagen type I (AteloCell, Japan) and 30 μg/mL fibronectin from human plasma (Sigma-Aldrich) in PBS buffered with 15 mM HEPES solution (Gibco) for 1-2 hours at 37°C. If not used directly, coated chips were stored at 4°C and pre-activated before use by incubation for 30 minutes with the same ECM solution at 37°C. Endothelial cells were cultured overnight at 37°C and 5% CO2 in T-75 cell culture flasks, detached with 0.05% Trypsin, concentrated to 5-10 million cells/mL, and seeded on the bottom face of the PDMS membrane. The chip was then incubated for a short period at 37 °C to allow the endothelial cells to spread and subsequently seeded with epithelial cells. Epithelial cells were seeded directly from cryopreserved vials received from the supplier, each vial of 0.5 million cells could be used to seed two chips. To increase reproducibility, vials were pooled and used to seed multiple chips. The chip was incubated overnight with complete epithelial and endothelial media in the epithelial and endothelial channels, respectively, under static conditions. The next day, the chip was washed and a reduced medium for the air-liquid interface (ALI) was flowed through the vascular channel using syringe pumps (Aladdin-220, Word Precision Instruments) at 60 μL/hour as described ^61^. The composition of the ALI media used was as described in ^61^ but with an FBS concentration of 5%. The epithelial face was incubated with epithelial base medium with 1 μM dexamethasone (Sigma Aldrich) without FBS supplementation to promote tight junction formation and surfactant expression as reported in previous lung-on-chip studies^61,62^. Flow was maintained over 2-3 days with daily replacement of the medium on the epithelial face (with dexamethasone supplementation). At the end of this period, in LoCs reconstituted with macrophages, CD14+ macrophages differentiated for 7 days in M-CSF (described above) were detached from the petri dish using 2 mM ethylenediaminetetraacetic acid (EDTA, Sigma Aldrich) in PBS at 4 °C as well as mechanical scraping, centrifuged at 300 *g* for 5 minutes, and resuspended in a small volume of epithelial cell media without dexamethasone. This solution containing macrophages was introduced onto the epithelial face and incubated for 30 minutes at 37 °C and 5% CO2 to allow macrophages to attach to the epithelial cells. Medium on the epithelial face was then removed and the chip was maintained overnight at an air-liquid interface (ALI). Chips were controlled to ensure that they successfully maintained the ALI overnight and were then transferred to the biosafety level 3 (BSL-3) facility for SARS-COV-2 infection. No antibiotics were used in any of the cell culture media for setting up the LoC model.

### Tocilizumab perfusion in the LoC model

Tocilizumab/Rho-Actemra (Roche, 20 mg/mL in saline) for intravenous administration were a kind gift from Prof Thomas Hugle at the University Hospital in Lausanne, Switzerland (CHUV). We simulated a single dose of 8 mg/kg given to a person of bodyweight 70 kg and a blood volume of 5 litres to obtain a concentration of 56 μg/mL in the cell culture media. Tocilizumab perfusion was started immediately after infection and was flowed through the vascular channel of infected chips for a period of 2 days post infection.

### Characterization of infection in LoCs

Uninfected LoCs were maintained at an ALI for either 1 day (for total RNA extraction (two LoCs without macrophages) or for up to 3 days for RNAscope characterization (one LoC each were reconstituted with and without macrophages) after addition of macrophages, during which time ALI medium was flowed through the endothelial channel at 60 μL/hour. In the case of the former, total RNA from the epithelial and endothelial layers was extracted separately by passing 350 μL of the lysis buffer reconstituted as per the manufacturer’s instructions (RNeasy Plus Micro Kit, Qiagen) through easy layer. Complete detachment of the cells in each layer was verified via optical microscopy. RNA was eluted in 25 μL of DEPC-treated nuclease free water and used for subsequent qRT-PCR reactions.

In the case of LoCs processed for RNAscope, the chips were fixed for 1 hour using a fresh solution of 4% paraformaldehyde in phosphate buffered saline (PBS), and subsequently dehydrated in a sequence of washes with 50% EtOH, 70% EtOH, and 100% EtOH. The chips were then stored in 100% EtOH at −20 °C until processed for RNAscope at the Histology Cor Facility at EPFL.

### Immunofluorescence characterization

For immunofluorescence labelling, the fixed LoCs were incubated with a blocking solution of 2% Bovine Serum Albumin (BSA) in PBS (‘blocking buffer’), followed by overnight incubation with the primary antibody in the blocking buffer at 4°C. A list of primary antibodies and concentrations used is included in Table S2. In addition, the infected LoC shown in Fig. 4 E-G was immunostained with Rabbit anti-SARS-CoV spike antisera ^63^, which was a kind gift of Prof Carolyn Machamer at John Hopkins University. The following day the chip was washed extensively with fresh blocking buffer before inoculation with the secondary antibody at a concentration of 1:300 for 1 hour at room temperature. The secondary antibodies used were: Donkey anti-mouse Alexa Fluor 488 (A21206, Thermo Fisher), or Donkey anti-mouse Alexa Fluor 568 (A10037, Thermo Fisher), or Donkey anti-mouse Alexa Fluor 647 (A31573, Thermo Fisher), and were chosen to complement the fluorophores already assigned to RNAscope labelling. F-actin on both the epithelial and endothelial face was stained using Sir-Actin dye in the far-red (Spherochrome) at 1 μM for 30 minutes concurrently with Hoechst staining, as described above. Confocal images were obtained on a Leica SP8 microscope in the inverted optical configuration at the EPFL BIOP core facility.

### Infection of the LoC with SARS-CoV-2

LoCs for infection were assembled into a sealed Tupperware box modified to contain inlets and outlets for the flow of medium through the vascular channel and was maintained throughout the course of the experiment by use of a syringe pump. On the day of the experiment, an aliquot of virus-containing supernatant was thawed from the stock at −80 °C, and diluted approximately 35-fold in epithelial cell media without FBS to generate the inoculum that corresponded to an infectious dose of 200 – 300 plaque forming units (PFU) in a volume of 30 μL. 30 μL of this solution was then added to the apical channel of each LoC, and the LoC was incubated for an hour at 37°C and 5% CO2 to enable infection. Thereafter, the medium in the apical channel was removed. The LoC was returned to air-liquid interface and the inlets of the infected chip were sealed with solid pins as a safety precaution. The media flowing through the vascular channel of each LoC was collected in a separate, sealed 15 mL Falcon tube, and each tube was emptied daily, and the effluent stored at −80 °C for further processing. At this time, the metal pins sealing the apical channel of each chip were removed, and 30 μL of epithelial cell media without FBS was briefly added to the channel before being withdrawn and collected as the ‘apical wash’. The metal pins were reinserted, and the LoCs returned to the cell culture incubator. Both the vascular effluent and the apical wash from each chip were collected for each day post infection.

Infection was terminated at specified time points either by treatment with the lysis buffer for RNA extraction (one LoC reconstituted without macrophages) or by fixing with freshly prepared 4% paraformaldehyde for a period of 1 hour (all other infected LoCs) for RNAscope assay preparation. The chips were then dehydrated as described above and immersed in 100 % EtOH before removal from the BSL-3 facility.

### RNAscope Assay

RNAscope Multiplex Fluorescent V2 assay (Bio-techne, Cat. No. 323110) was performed according to manufacturer’s protocol directly on chips. They were hybridized with the control Hs 3plex positive control (Bio-techne, Cat. No. 320861) or a combination of the following probes V-nCoV2019-orf1ab-sense (Bio-techne, Cat. 859151), Hs-ACE2-C2 (Bio-techne, Cat. 848151-C2), V-nCoV2019-S-C3 (Bio-techne, Cat. 848561-C3), HS-Il1b-C1 (Bio-techne, Cat. 310361), and Hs-Il6-C2 (Bio-techne, Cat. No. 310371-C2) simultaneously at 40°C for 2 hours. The different channels were revealed with TSA Opal570 (Akoya Biosciences, Cat. No. FP1488001KT), TSA Opal650 (Akoya Biosciences, Cat. No. FP1488001KT) and TSA Opal690 (Akoya Biosciences, Cat. No. FP1497001KT). Cells were counterstained with DAPI and mounted with Prolong Gold Antifade Mountant (Thermo Fisher, P36930).

### ELISA assays

Frozen vascular effluent from infected chips was thawed, incubated at 58 °C for 30 minutes to ensure sterilization, and then removed from the BSL-3 for further processing. Levels of human IL-6 were determined by enzyme-linked immunosorbent assays (BD Biosciences) as per the manufacturers’ instructions.

### Confocal imaging and Image Analysis

Infected and control LoCs were imaged using a Leica SP8 confocal microscope with a white light laser. For chips processed via RNAscope and labelled with Opal 650 and Opal 690, the excitation and emission windows were carefully chosen to minimise overlap of signal. LoCs were imaged with a 25x water immersion objective (NA=0.95, Leica), with standard settings across chips labelled the same way. Z stacks were subsequently deconvolved using the Huygens Deconvolution Software (Scientific Volume Imaging) and 3D views were rendered using the FIJI ClearVolume plugin^64^. Custom-written software in MATLAB was used to segment and identify the 3D volume, mean intensity, and number of RNA dots in each field of view using the nestedSortStruct algorithm for MATLAB written by the Hughey Lab (https://www.github.com/hugheylab/nestedSortStruct, GitHub). Statistical analysis was performed using Origin 9.2 (OriginLabs) and P-values were calculated using a Kruskal-Wallis one-way ANOVA test, with the null hypothesis that the medians of each population were equal.

## Data availability statement

The datasets generated during and/or analysed during the current study are available from the corresponding authors on reasonable request. Software code used to analyse these datasets will be uploaded to Zenodo prior to publication

