## Supplementary Material for "Rapid endothelial infection, endothelialitis and vascular damage characterise SARS-CoV-2 infection in a human lung-on-chip model"

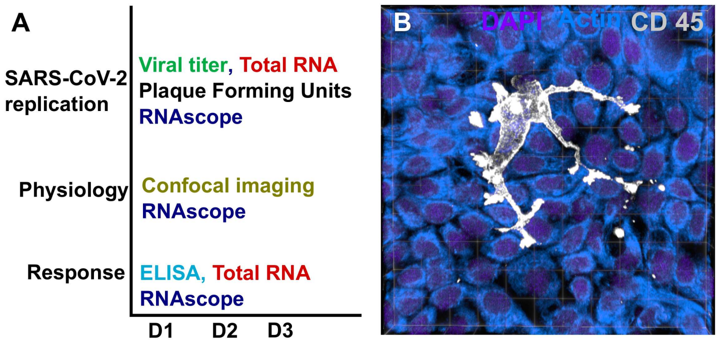


**Fig. S1.**

(**A**) An overview of the techniques used to characterize viral replication, physiological changes, and the cellular responses in the LoC over 3 days post infection. (**B**) 3D view of the epithelial layer of a an uninfected control LoC reconstituted with CD14+ macrophages added to the epithelial side. Macrophages are identified via an anti-CD45 antibody (grey), actin and nuclear labelling are labelled in azure and electric indigo LUTs respectively.

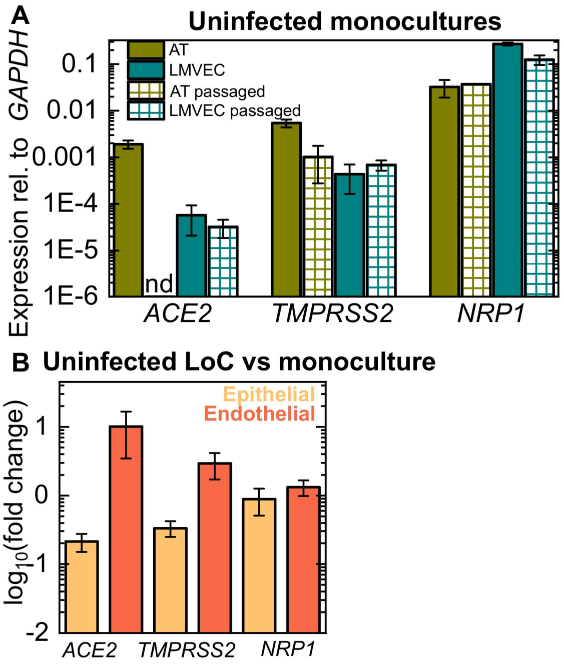


**Fig. S2.**

(**A**) Plots of expression of the cell receptors *ACE2* and *NRP1*, and the protease *TMPRSS2* relative to *GAPDH* expression for alveolar epithelial cells (ATs) obtained from a commercial supplier at passage 3, alveolar epithelial cells post-passage in the lab ('AT passaged'), freshly isolated lung microvascular endothelial cells from a commercial supplier ('LMVEC') and lung microvascular endothelial cells post-passage ('LMVEC passaged'). 'nd' refers to not detected. (**B**) Plot of the fold change in expression of viral entry factors in cells from the epithelial and endothelial layers of n=3 uninfected control LoC vs monocultures.

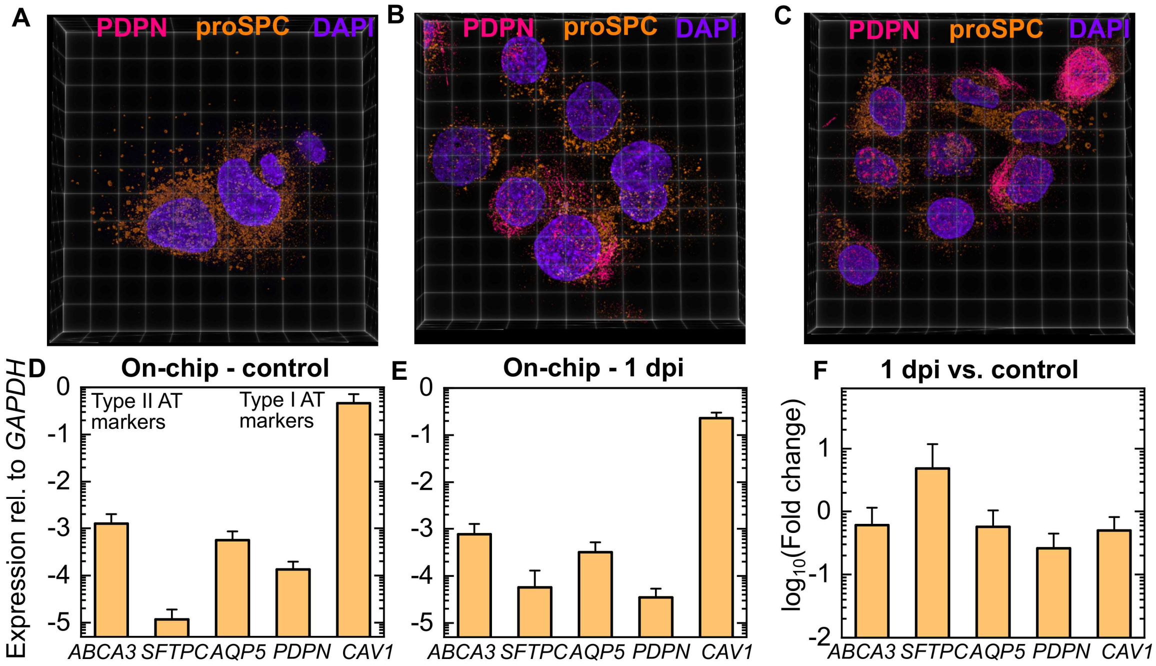


**Fig. S3.**

(**A-C**) 3D views of individual alveolar epithelial cells at passage 3. Lamellar bodies characteristic of type II alveolar epithelial cells are identified via an anti pro-SPC antibody (shown in amber), type I alveolar epithelial cells were identified via an anti Podoplanin antibody (*PDPN*, shown in pink). Nuclear labelling is shown in electric indigo. (**D, E**) Plots of the expression of type II AT markers (*ABCA3*, *SFTPC*) and type I AT markers (*AQP5*, *PDPN*, *CAV1*) relative to *GAPDH* expression from cells in the epithelial layer of uninfected controls (n=3, **D**) and infected LoCs at 1 dpi (n=3, **E**). (**F**) Plots of the fold change in AT markers from the epithelial layers of the LoCs in (**D**) and (**E**). In all plots, the bar represents the mean and the error bars the standard deviation.

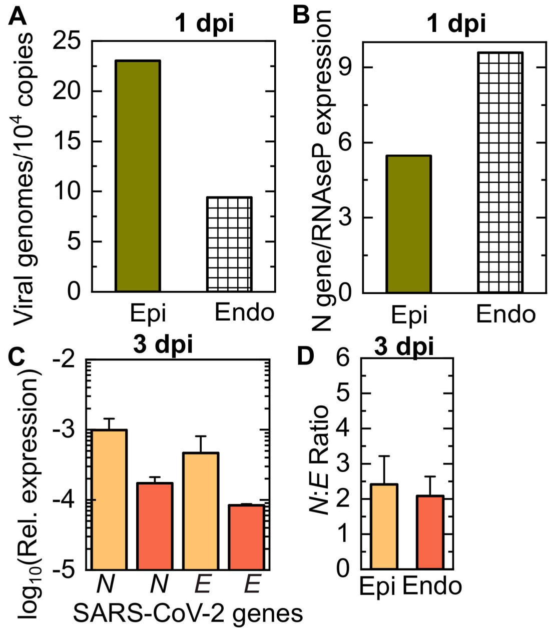


**Fig. S4.**

(**A**) Quantification of the numbers of viral genomes detected in the total RNA obtained from the apical and vascular channels respectively of an infected LoC without macrophages at 1 dpi. (**B**) Intracellular viral RNA levels at this timepoint relative to levels of the eukaryotic housekeeping gene *RNaseP. (***C**) Quantification of the intracellular levels of transcripts for the *SARS-CoV-2 N* and *SARS-CoV-2 E* genes in the epithelial and endothelial layers of infected LoCs at 3 dpi (n=2) relative to expression of the eukaryotic housekeeping gene GAPDH. (**D**) SARS-CoV-2 N : SARS-CoV-2 E ratio in the epithelial and endothelial layers of the infected LoCs shown in (**C**). The bars represent the mean value and the error bars represent the standard deviation.

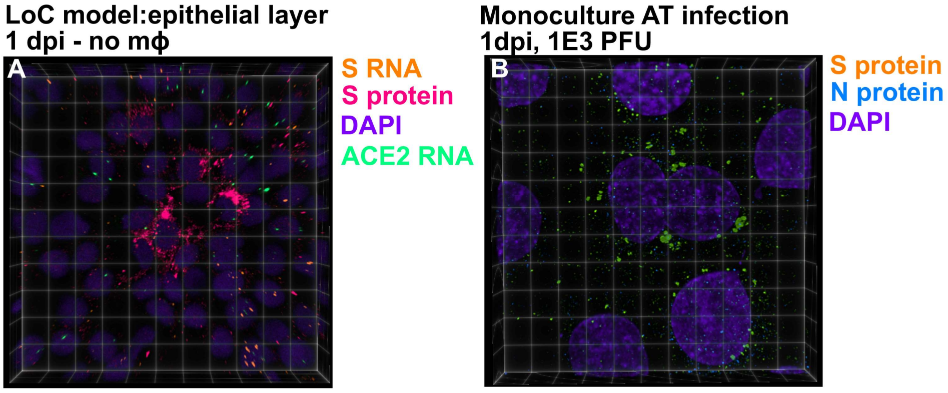


**Fig. S5.**

(**A**) 3D view of a 232 x 232 μm$^2$ field of view of the epithelial layer of an infected LoC at 1 dpi. Three cells with productive amplification of virions can be identified by the formation of clusters of spike protein (identified via antibody labelling and shown in bright pink LUT) localized in the cytoplasm surrounding the nucleus. S RNA and *ACE2* mRNA identified via an RNAscope assay and nuclear labelling are shown in amber, spring green, and electric indigo LUTs respectively. (**B**) 3D view of a 61.4 x 61.4 μm$^2$ field of view of an alveolar epithelial cell monolayer at 1 dpi. Cells with productive amplification of virions can be identified by the formation of clusters of spike protein (identified via antibody labelling and shown in amber LUT) localized in the cytoplasm surrounding the nucleus. SARS-CoV-2 nucleocapsid (N) protein and nuclear labelling are shown in amber and electric indigo LUTs respectively.

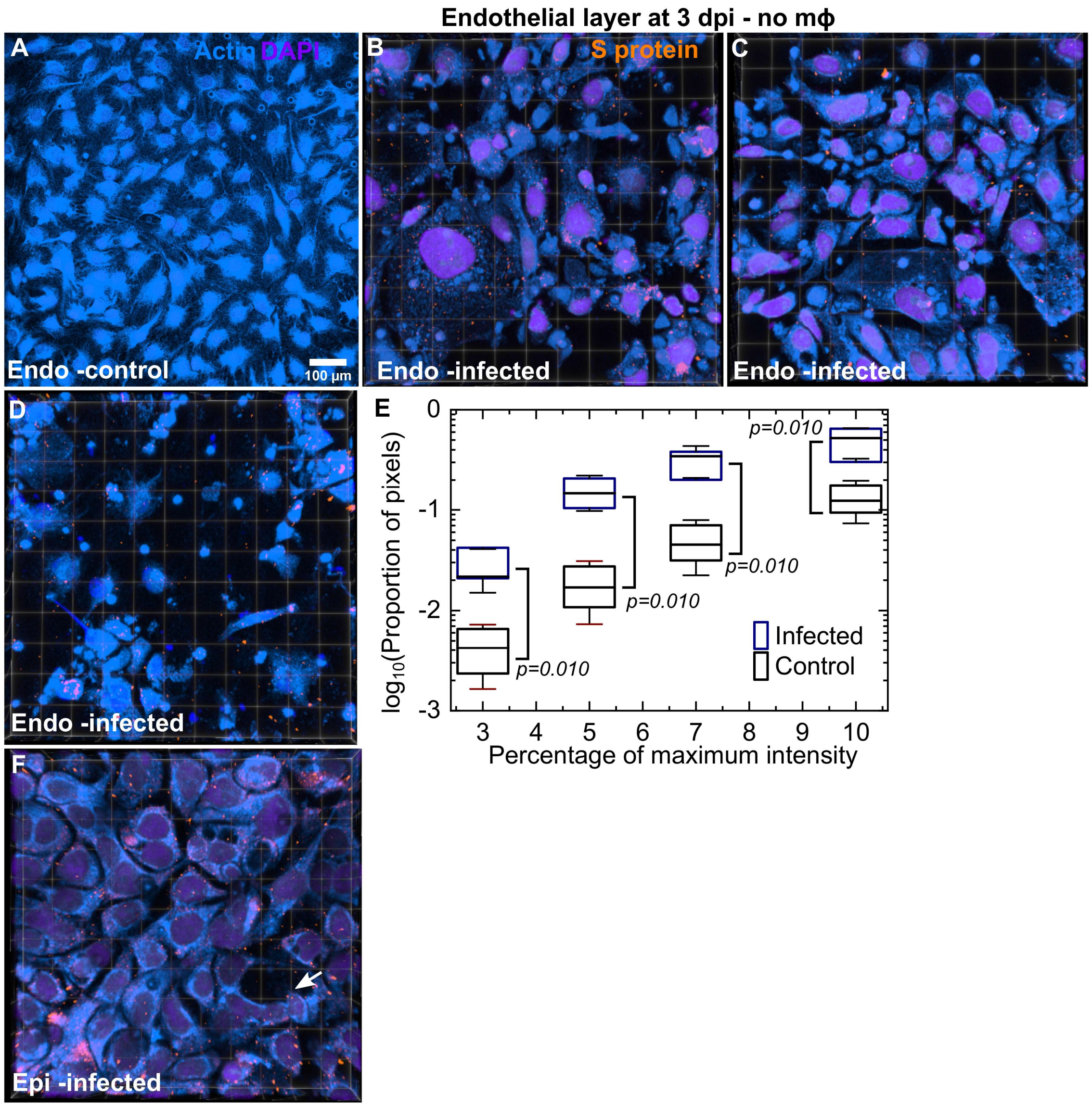


**Fig. S6**

(**A**) A maximum intensity projection of a 465 x 465 µm$^2$ field-of-view of the endothelial layer in an uninfected chip at 3 days post air-liquid interface shows intact vasculature. (**B**-**D**) Additional examples of endothelial damage – 3D views of 232 x 232 µm$^2$ fields-of-view from LoCs infected without macrophages at 3 dpi. S protein identified via antibody labelling, actin and nuclear labelling are shown in amber, azure and electric indigo LUTs respectively. Many areas with little or no actin staining are observed. A plot of the proportion of pixels (shown on a logarithmic scale) with regions of low F-actin intensity identified via cut-off thresholds defined as a percentage of the maximum intensity in 4-5 fields of view each for control and infected LoCs. P-values are calculated using a one-way Kruskal-Wallis ANOVA test. (**F**) An additional example of a 3D view of a 155 x 155 µm$^2$ field of view of the epithelial layer from the same LoC shown in (**B**-**D**). The integrity of the layer is better maintained, cell loss is indicated by a white arrow.

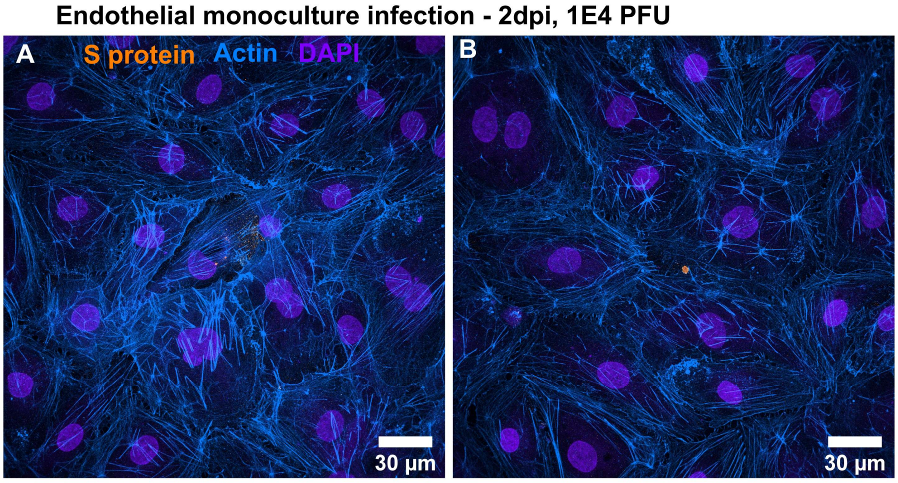


**Fig. S7**

(**A**, **B**) Representative images of endothelial cell infection in a 24 well plate, 2 days post infection with an infectious dose of 1E4 plaque forming units (PFU). An intact monolayer and few signs of SARS-CoV-2 infection are seen. S protein is detected via immunofluorescence and colored amber, Actin is colored azure, and nuclear staining is colored indigo.

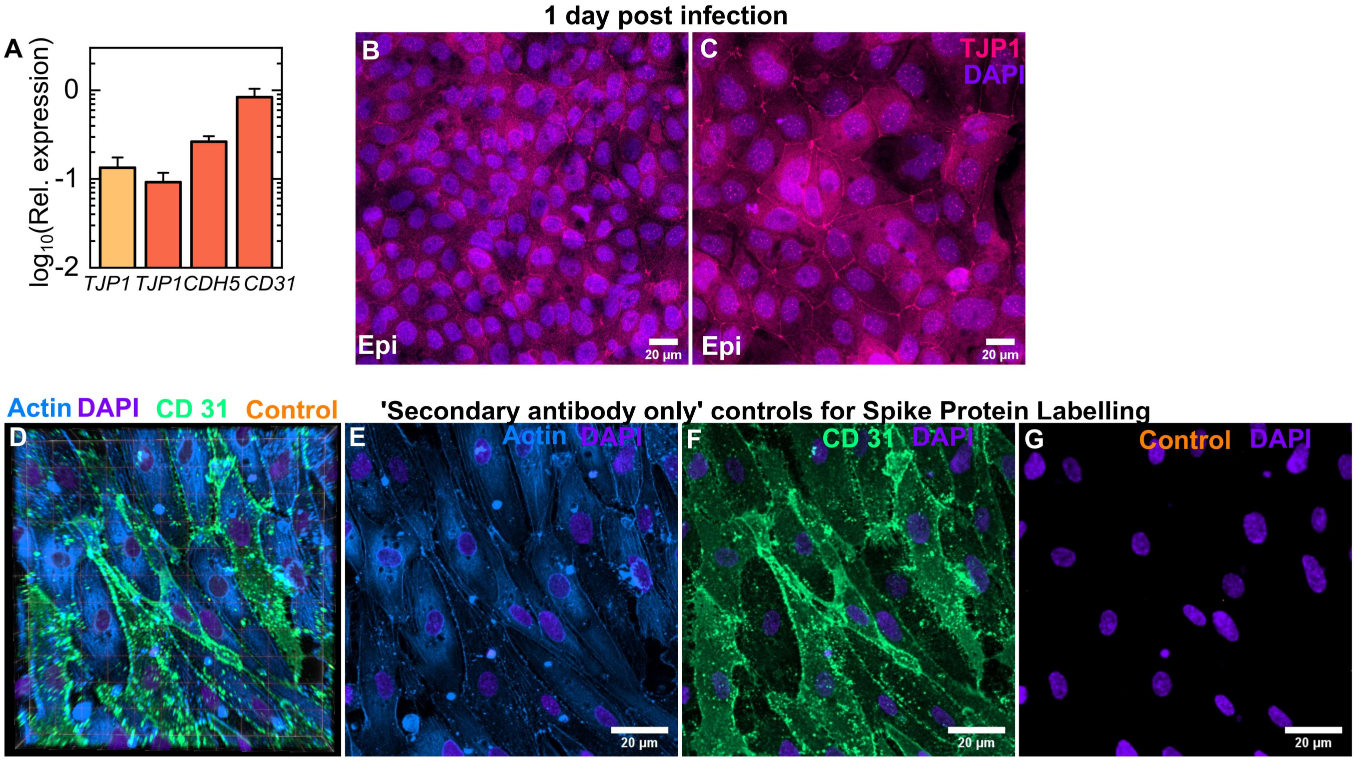


**Fig. S8**

(**A**) Plot of the expression of the tight junction markers PECAM-1 (*CD31*) and VE-Cadherin (*CDH5*) in cells from the endothelial layer and of ZO-1 (*TJP1*) in cells from both the endothelial and epithelial layers of uninfected controls (n=3) relative to *GAPDH* expression. The bars represent the mean, and the error bars represent the standard deviation. (**B**, **C**) Maximum intensity projection of two additional fields of view from the epithelial layer of an infected LoC reconstituted without macrophages at 1 dpi. TJP1 identified via antibody labelling and nuclear labelling are shown in bright pink and electric indigo LUTs respectively. (**D**-**G**) Secondary antibody only controls for spike protein labelling. (**D**) 3 D view of a 116.36 x 116.36 mm2 field of view of the endothelial layer of an uninfected control LoC. Actin, CD31, secondary antibody only ('control') and nuclear labelling are indicated in azure, spring green, amber, and electric indigo LUTs respectively. (**E**-**G**) Maximum intensity projections of actin (**E**), CD31 (**F**) and control staining (**G**) for the same field of view. Nuclear labelling is shown in all panels.

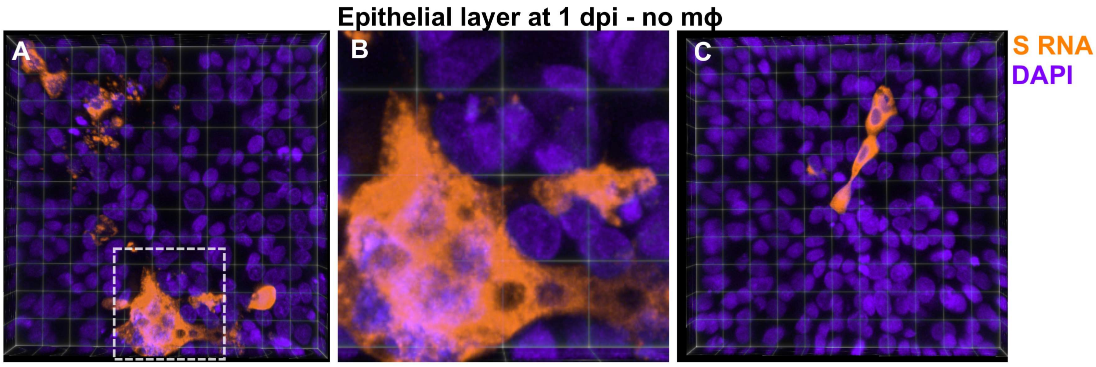


**Fig. S9.**

(**A**, **C**) 3D views of two representative 232 x 232 $\mu m^2$ fields of view of the epithelial layer of an LoC reconstituted without macrophages at 1 dpi. S RNA is identified by RNAscope assay and false colored amber, and nuclear labelling is false colored indigo. Each field of view shows examples of heavily infected cells. (**B**) Zoom corresponding to the area marked by the white box in (**A**), a collection of heavily infected cells is visible together with syncytia formation.

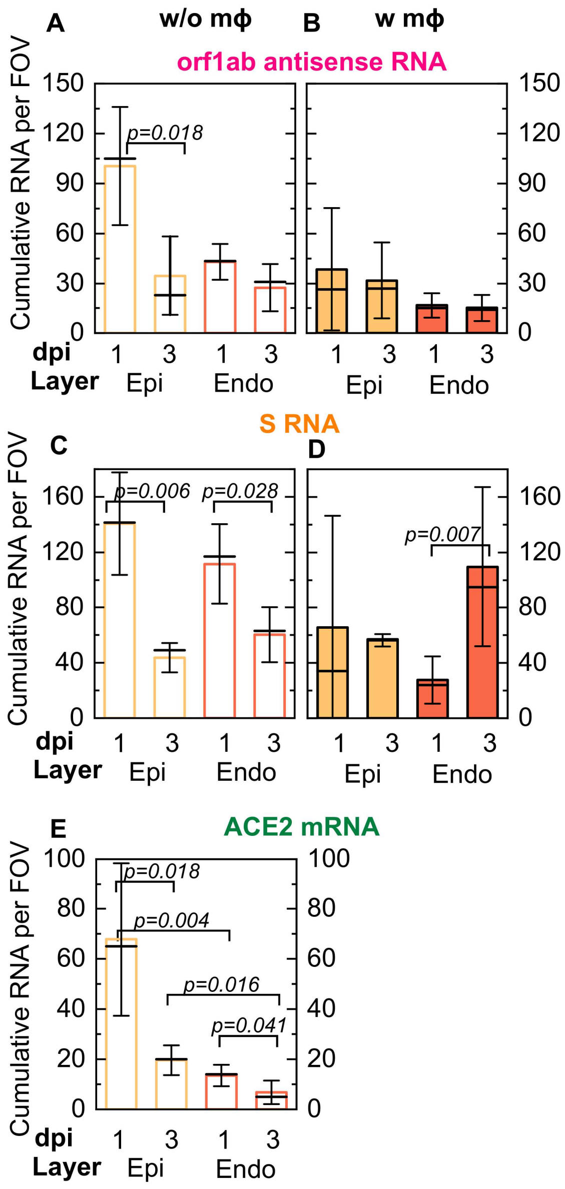


**Fig. S10**

Quantification of viral antisense RNA (**A, B**), viral genomic RNA (**C, D**) and *ACE2* mRNA from pairs of otherwise identical LoCs reconstituted without (**A, C, E**) and with macrophages (**B, D**) and analyzed at 1 and 3 dpi. Plots show the cumulative number of spots imaged per field of view from 4-6 field of views detected using RNAscope and confocal imaging using identical imaging conditions for all chips. Bars represent the mean value, the solid line represents the median, and error bars represent the standard deviation. (**E**) SARS-CoV-2 infection reduces *ACE2* expression per field of view in both epithelial cells (p=0.018) and endothelial cells by 3 dpi (p=0.041) in LoCs reconstituted without macrophages. P-values are calculated using a Kruskal-Wallis One-Way ANOVA Test.

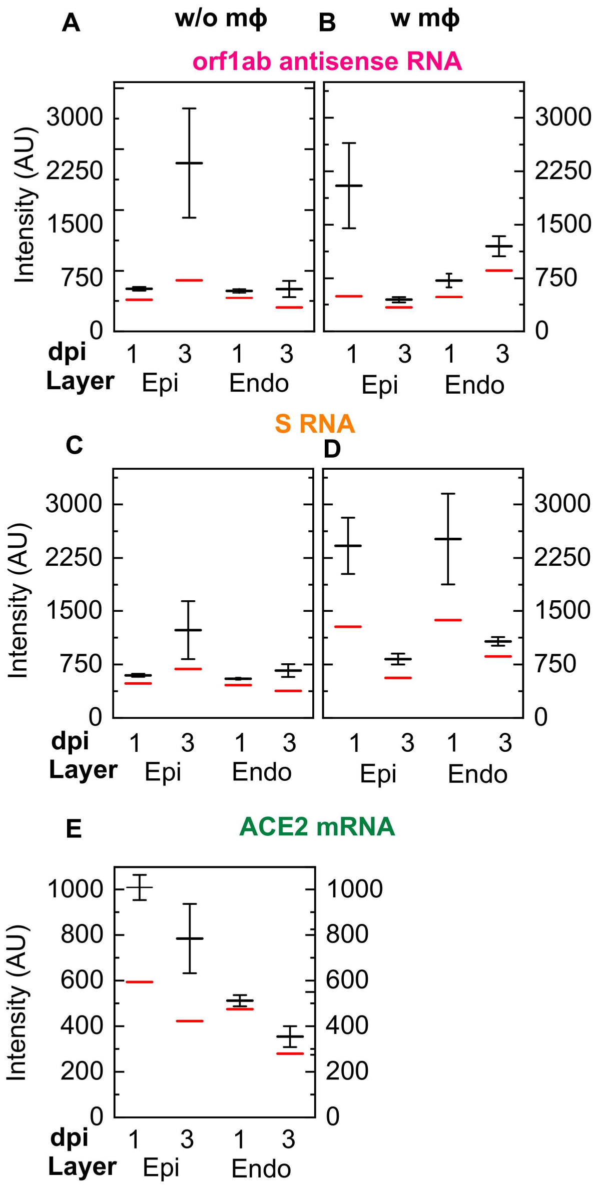


**Fig. S11**

Quantification of viral antisense RNA (**A, B**), viral genomic RNA (**C, D**) and *ACE2* mRNA from pairs of otherwise identical LoCs reconstituted without (**A, C, E**) and with macrophages (**B, D**) and analyzed at 1 and 3 dpi. Plots show the mean (solid black line) and median (solid red line) of the intensity of each spot detected using RNAscope and confocal imaging using identical imaging conditions for all chips. The error bars represent the standard deviation.

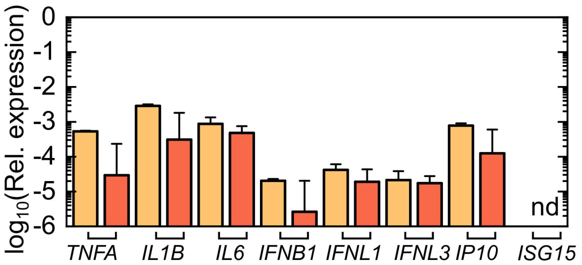


**Fig. S12**

Plot of the expression of the NF-KB related pro-inflammatory cytokines (*TNFA*, *IL1B*, *IL6*), interferon genes (*IFNB1*. *IFNL1*, and *IFNL3*) and interferon stimulated genes (*IP10*, *ISG15*) in cells from the epithelial and endothelial layers of uninfected controls (n=3). ‘nd’ refers to not detected. The bars represent the mean, error bars represent the standard deviation.

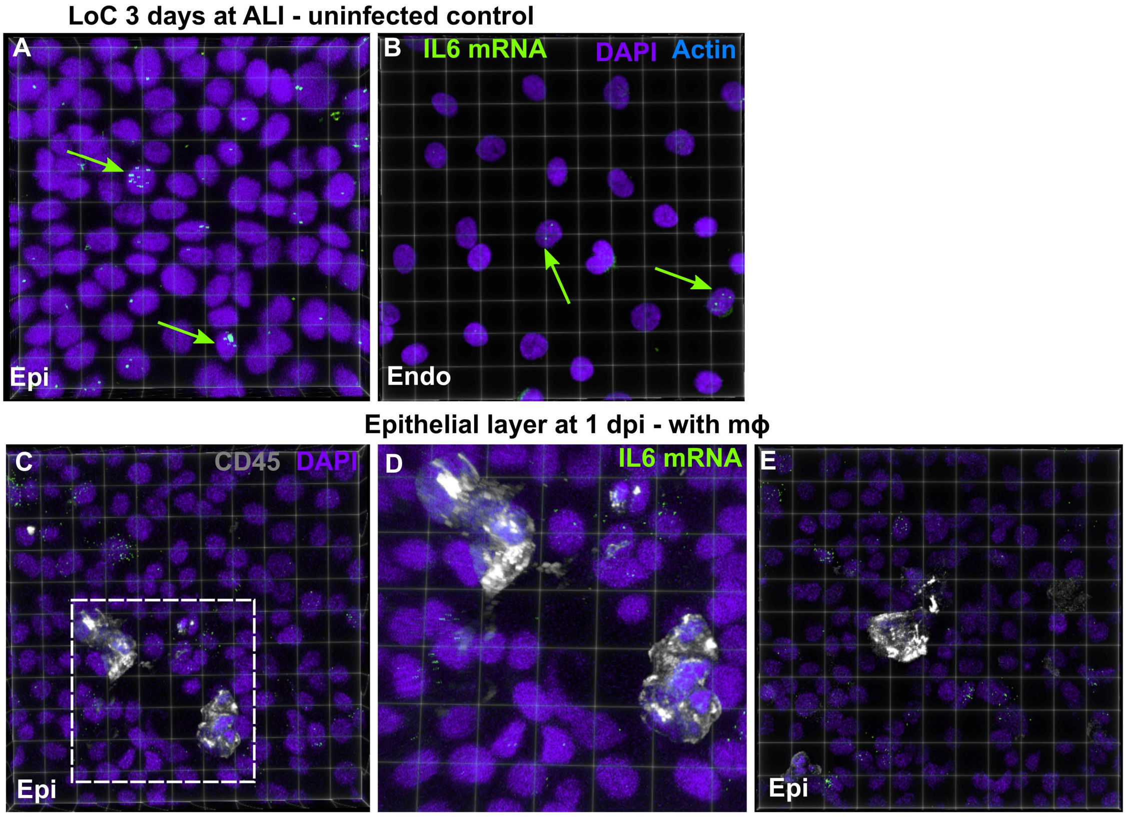


**Fig. S13**

(**A**, **B**) 3D views of representative 155 x 155 μm$^2$ fields of view of the epithelial layer and endothelial layer corresponding to the images Fig. 1B and 1C. *IL6* mRNA is identified by RNAscope assay and false colored chartreuse (yellow arrows) , and nuclear labelling is false colored indigo. Yellow arrows indicate *IL6* mRNA co-localised with the nucleus. (**C, E**) 3D views of two 232 x 232 μm$^2$ fields of view of the epithelial layer of an infected LoC reconstituted with macrophages at 1 dpi. (**D**) Zoom corresponding to the area marked by the white box in (**C**), macrophages are labelled via an anti-CD45 antibody (gray).

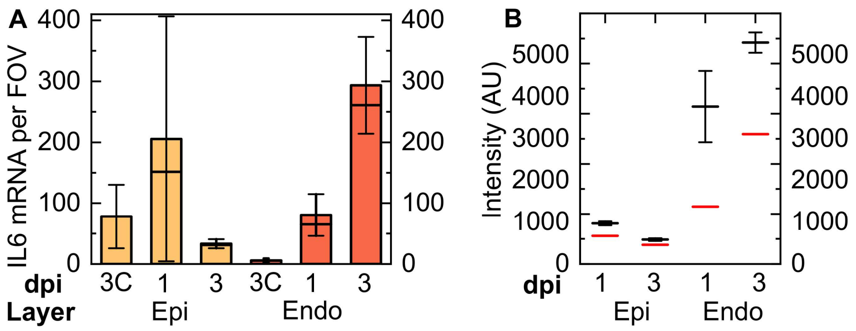


**Fig. S14.**

(**A**) Quantification of *IL6* expression in epithelial and endothelial cells from a pair of otherwise identical LoCs analyzed at 1 and 3 dpi, respectively. Plots show the total number of spots in 4-6 fields of view detected using RNAscope assay using identical imaging conditions for all chips. Bars represent the mean value, the solid line represents the median, and error bars represent the standard deviation. Data from control uninfected LoCs corresponding to the 3 dpi timepoint is labelled ‘3C’. (**B**) Plots show the mean (solid black line) and median (solid red line) of the intensity of each spot detected using RNAscope and confocal imaging using identical imaging conditions for all chips. The error bars represent the standard deviation.

| **Sequence (5' - 3')** | **Primer** |
|---|---|
| TAC GGT GTG GCA CCG CTC AAT G | *AQP5* forward |
| AGT CAG TGG AGG CGA AGA TGC A | *AQP5* reverse |
| CCA AGG AGA TCG ACC TGG TCA A | *CAV1* forward |
| GCC GTC AAA ACT GTG TGT CCC T | *CAV1* reverse |
| GTG CCG AAG ATG ATG TGG TGA C | *PDPN* forward |
| GGA CTG TGC TTT CTG AAG TTG GC | *PDPN* reverse |
| GTC CTC ATC GTC GTG GTG ATT G | *SFTPC* forward |
| AGA AGG TGG CAG TGG TAA CCA G | *SFTPC* reverse |
| CTT GAC AGT CGC AGA GCA CCT T | *ABCA3* forward |
| CTC CGT GAG TTC CAC TTG TCC T | *ABCA3* reverse |
| GTC TCC TCT GAC TTC AAC AGC G | *GAPDH* forward |
| ACC ACC CTG TTG CTG TAG CCA A | *GAPDH* reverse |
| TCC ATT GGT CTT CTG TCA CCC G | *ACE2* forward |
| AGA CCA TCC ACC TCC ACT TCT C | *ACE2* reverse |
| CCT CTA ACT GGT GTG ATG GCG T | *TMPRSS2* forward |
| TGC CAG GAC TTC CTC TGA GAT G | *TMPRSS2* reverse |
| AAC AAC GGC TCG GAC TGG AAG A | *NRP1* forward |
| GGT AGA TCC TGA TGA ATC GCG TG | *NRP1* reverse |
| CAT TAC CTG AAG GCC AAG GA | *IFNB1* forward |
| CAG CAT CTG CTG GTT GAA GA | *IFNB1* reverse |
| AAC TGG GAA GGG CTG CCA CAT T | *INFL1* forward |
| GGA AGA CAG GAG AGC TGC AAC T | *IFNL1* reverse |
| TCG CTT CTG CTG AAG GAC TGC A | *IFNL3* forward |
| CCT CCA GAA CCT TCA GCG TCA G | *IFNL3* reverse |
| TGA GGT ACA GGC CCT CTG AT | *TNFA* forward |
| CCC GAG TGA CAA GCC TGT AG | *TNFA* reverse |
| CCA CCT CCA GGG ACA GGA TA | *IL1B* forward |
| AAC ACG CAG GAC AGG TAC AG | *IL1B* reverse |
| ATT TGC CGA AGA GCCC TCA G | *IL6* forward |
| CCC CTG ACC CAA CCA CAA AT | *IL6* reverse |
| TGA TGG CCT TCG ATT CTG GA | *IP10* forward |
| AGT GGC ATT CAA GGA GTA CC | *IP10* reverse |
| CAG CCA TGG GCT GGG AC | *ISG15* forward |
| GCC GAT CTT CTG GGT GAT CT | *ISG15* reverse |
| AAC AGC GAC TGC ACG TTG AAG G | *ADAM17* forward |
| CTG TGC AGT AGG ACA CGC CTT T | *ADAM17* reverse |
| GAC TGT GCA CTT GCT GGT GGA T | *IL6R* forward |
| ACT TCC TCA CCA AGA GCA CAG C | *IL6R* reverse |
| CAG AGT TCA CAC CTT ACC TGG AG | *F3* forward |
| GTT GTT CCT TCT GAC TAA AGT CCG | *F3* reverse |

| | |
|---|---|
| CAG CTC AAT GCT GTG AAT AAC TCC | *TFPI* forward |
| TCT GCT GGA GTG AGA CAC CAT G | *TFPI* reverse |
| AAC GAC CTC TGC GAG CAC TTC T | *THBD* forward |
| CCA GTA TGC AGT CAT CCA CGT C | *THBD* reverse |
| CTC ATC AGC CAC TGG AAA GGC A | *SERPINE1* forward |
| GAC TCG TGA AGT CAG CCT GAA AC | *SERPINE1* reverse |
| CCT TGA ATC CCA GTG ACC CTG A | *VWF* forward |
| GGT TCC GAG ATG TCC TCC ACA T | *VWF* reverse |
| AAG TGG AGT CCA GCC GCA TAT C | *CD31* forward |
| ATG GAG CAG GAC AGG TTC AGT C | *CD31* reverse |
| GAA GCC TCT GAT TGG CAC AGT G | *CDH5* forward |
| TTT TGT GAC TCG GAA GAA CTG GC | *CDH5* reverse |
| GTC CAG AAT CTC GGA AAA GTG CC | *TJP1* forward |
| CTT TCA GCG CAC CAT ACC AAC C | *TJP1* reverse |
| CAA TGC TGC AAT CGT GCT AC | *SARS-CoV-2 N* forward |
| GTT GCG ACT ACG TGA TGA GG | *SARS-CoV-2 N* reverse |
| TCG TTT CGG AAG AGA CAG GT | *SARS-CoV-2 E* forward |
| GCG CAG TAA GGA TGG CTA GT | *SARS-CoV-2 E* reverse |

**Table S1.**
Primers used for qRT-PCR characterization of gene expression in this study.

| Antibody | Supplier | Catologue Number | Staining Concentration |
|---|---|---|---|
| anti-mouse proSPC (Rabbit polyclonal) | Abcam | Cat#: ab40879 | IF (1:100) |
| anti- human Podoplanin / gp36 (Mouse monoclonal [18H5] | Abcam | Cat#: ab10288; | IF (1:100) |
| anti-human von Willebrand Factor conjugated to Alexa 647 (Rabbit monoclonal [EPSISR15]) | Abcam | Cat#: ab195029 | IF (1:100) |
| anti-human ZO-1 (Rabbit polyclonal) | Abcam | Cat#: ab216880 | IF (1:100) |
| anti-human CD 31 (Mouse monoclonal [P2B1]) | Abcam | Cat#: ab24590 | IF (1:100) |
| anti-human CD 5 (Mouse monoclonal [MEM-28]) | Abcam | Cat#:ab8216 | IF (1:100) |
| anti-SARS-COV-2 Spike protein (Mouse monoclonal [1A9]) | Genetex | Cat#: GTX632604 | IF (1:750) |
| anti-SARS-COV-2 Nucleocapsid protein (Mouse monoclonal ([6H3]) | Genetex | Cat#:GTX632269 | IF (1:750) |

**Table S2.**
Commercially available primary antibodies used in this study.
